# Autoantibody origins in lupus and in relapse post CAR-T therapy

**DOI:** 10.1101/2025.10.20.683393

**Authors:** Amalie Grenov, Jongwon Yoon, Daniel M. Snell, Anna Mikolajczak, Hao Wang, Erdem Gürel, Ana Rodriguez Ronchel, Gülçin Yegen, Lydia Lee, Bahar Artim Esen, Anisur Rahman, Paul Maciocia, Carola G. Vinuesa

## Abstract

Anti-CD19 chimeric antigen receptor (CAR)-T therapy induces profound remissions in lupus by depleting B cells, challenging the longstanding view that treatment-resistant disease is sustained by long-lived plasma cells. Additionally, emerging relapses highlight the need to understand how pathogenic autoantibodies arise. Using molecular antibody tagging in mice with human monogenic lupus variants, we reveal that autoantibody-producing cell cohorts are long-lived but plasma cells are short-lived, requiring continuous replenishment from proliferating precursors. The spleen acts as a major plasma cell reservoir, with perivascular localization conserved in mice and lupus patients. Relapse after anti-CD19 CAR-T occurred through newly-generated B cells rather than treatment-resistant clones. Plasma cell depletion by anti-BCMA CAR-T failed to eliminate some precursors that become autoantibody-secreting. These findings demonstrate that continuous B cell-to-plasma cell differentiation, not intrinsic plasma cell longevity, sustains pathogenic antibody responses in lupus, supporting a potential benefit of adjunctive therapies after CAR-T, particularly in genetically predisposed patients.

## Introduction

B cells are increasingly recognized as key pathogenic drivers across a spectrum of autoimmune diseases - a view reinforced by the recent clinical success of B cell-depleting CAR-T cell therapies (Mougiakakos, Meyer, and Schett 2025). Although B cells exert their effects through multiple mechanisms, autoantibody production remains a central driver and amplifier of disease. Autoantibodies often precede clinical onset, serve as valuable diagnostic biomarkers, and correlate with disease severity (Hale, Rawlings, and Jackson 2018; Choi, Kim, and Craft 2012). In the rapidly evolving field of B cell-depleting therapies, understanding the origin, persistence, and anatomical niches of autoantibody-producing cells is crucial.

Long-lived plasma cells, which classically express little or no CD20 or CD19, have been considered the main reservoirs of pathogenic antibodies, particularly given the persistence of autoantibodies following CD20-targeted therapies like rituximab (Hale, Rawlings, and Jackson 2018). Yet, the profound and sustained autoantibody reduction following anti-CD19 CAR-T cell therapy challenges this paradigm (Mougiakakos, Meyer, and Schett 2025). A leading hypothesis to explain rituximab’s inferior efficacy is insufficient tissue penetration, resulting in incomplete depletion of CD20⁺ memory and tissue-resident B cells (Kamburova et al. 2013; Ramwadhdoebe et al. 2019). The effectiveness of anti-CD19 CAR-T cells suggests that either autoreactive plasma cells are uniformly short-lived, or that disease maintenance depends more critically on CD19^+^ cells. However, findings from lupus-prone mouse models have been mixed. In NZB/W mice, long-lived plasma cells have been reported in various tissues using indirect methods such as BrdU incorporation (Khodadadi et al. 2015; Starke et al. 2011; Taddeo et al. 2015), and sustained depletion of autoantibody-secreting cells required combination therapy with proteasome inhibitors and anti-CD20 antibodies to target both plasma cells and their precursors (Taddeo et al. 2015; Khodadadi et al. 2015). Nevertheless, autoantibody-secreting cells re-appeared within two weeks following depletion by proteasome inhibitors (Taddeo et al. 2015; Khodadadi et al. 2015). It is plausible that more precise approaches to track plasma cells and their precursors in vivo may yield different results.

Clinical observations mirror this complexity. Although still rare, cases of myositis and lupus nephritis relapses following anti-CD19 CAR-T cell therapy have recently been reported within two 30-patient cohorts (Müller et al. 2025; Floersh 2024). These reports highlight the need to determine whether relapsing autoimmunity is driven by residual, depletion-resistant plasma cells or their precursors, or by newly generated autoreactive B cells. This distinction has direct therapeutic implications as plasma cells express BCMA, and BCMA-targeted CAR-T constructs are in clinical trials. If relapse is primarily driven by newly formed autoreactive B cells, CD19^+^ B-cell–directed therapies may outperform anti-BCMA strategies. A related question is whether patients with strong genetic predisposition will relapse more readily following CAR-T therapy. Approximately 7-10% of juvenile SLE cases are attributable to highly penetrant rare genetic variants (Belot et al. 2020); these patients tend to experience a severe clinical course, and have yet to be included in CAR-T cell trials.

The preferred anatomical locations of pathogenic plasma cells in systemic autoimmunity also remain poorly defined, yet this has direct therapeutic consequences. While bone marrow niches are well known to support long-term plasma cell survival (Hargreaves et al. 2001; Nutt et al. 2015; Robinson et al. 2023), inflamed peripheral tissues including the spleen, skin, joints, and kidneys, can also serve as supportive microenvironments for plasma cells (Khodadadi et al. 2019).

Addressing all these questions in human patients remains technically and ethically challenging. Access to critical tissues is constrained and tools for tracing the fate of B cell subsets in humans are still lacking. This hampers our ability to determine the cellular origin of autoantibodies, including following CAR-T intervention.

To overcome these barriers, we combined mouse models that carry genetic variants found to cause lupus in children with immunoglobulin-tagging methods. Using these tools, we show that plasma cells are short-lived and abundant in the spleen — a finding corroborated in spleen samples from human SLE patients. We further demonstrate that anti-CD19 CAR-T therapy efficiently eliminates B cells and plasma cells in the spleen, and that consequently, relapse following treatment is not caused by depletion-resistant plasma cells, but rather by newly generated autoreactive B cells. These insights suggest that introducing therapies that prevent B cell activation or maturation soon after CAR-T cell decline may prevent relapse, particularly in patients with strong genetic predisposition.

## Results

### Tracking the cellular origin of autoantibodies [Figure 1]

Antinuclear antibody immunoassays offer a snapshot of the circulating autoantibodies produced by any antibody secreting cell (ASC) that arose at any point in life. However, defining the cellular and temporal origin of autoantibodies requires fate-mapping of B cells unto the antibodies they produce as well as mouse models that closely replicate human disease. To this end, we took advantage of K-tag antibody-tracking mice (Schiepers et al. 2023). In heterozygous K-tag (*Igk^Tag/WT^*) mice, approximately 50% of Igĸ^+^ B cells (gated as Igl^-^) carry a Flag-tag on the light chain (Figure 1A-C and Figure S1A-B), which irreversibly switches to a Strep-tag upon Cre-mediated recombination. In *Cd79a^ERT2cre^ Igk^Tag^* (*Cd79-Igk^Tag^*) mice, tamoxifen-induced Cre activation drives a complete switch from Flag to Strep expression across the *Igk^Tag^* B cell pool (Figure 1C, day 8). As illustrated in Figure 1A, this system enables time-stamping of B cells to track both retention of Strep-tagged cells present at tamoxifen induction and emergence of newly generated Flag-tagged B cells that develop in the bone marrow thereafter (Figure 1C days 30 and 52). Tagged antibodies from each population can be quantified simultaneously by ELISA.

**Figure 1:**
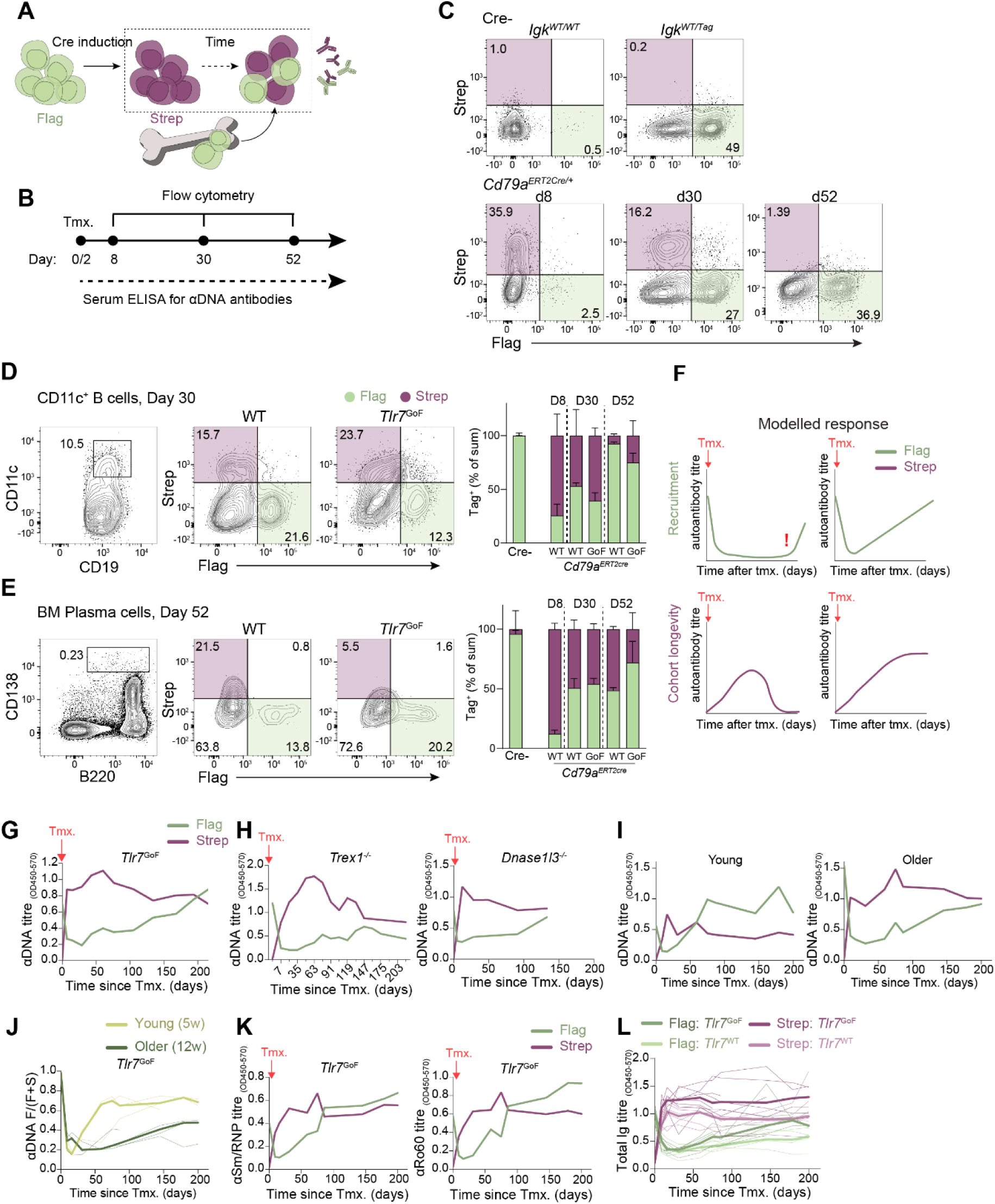
Autoreactive B cells are recruited continuously and give rise to long-lived cohorts of autoantibody producing cells. **(A)** Schematic showing tamoxifen-induced Flag-(green) to Strep-(purple) tag-switching in a pre-existing B cell cohort and replenishment over time by Flag-labelled (green) newly produced B cells in *Cd79a-Igk^Tag^* mice. **(B)** Timing of tamoxifen administration and flow cytometry analysis. **(C)** Flow cytometry plots showing Flag-and Strep-tag expression in mice with and without *Igk^Tag^* and *Cd79a^ERT2cre^* alleles. Bottom row shows consecutive time points after tamoxifen treatment. **(D-E)** Gating strategy for CD11c^+^ ABCs (D) and bone marrow plasma cells (E) (left) and surface expression of Flag- and Strep-tag 30 or 52 days after tamoxifen in WT and *Tlr7*^GoF^ mice. Relative contribution of Flag^+^ and Strep^+^ cells to total Igκ-tag^+^ cells quantified over time (right panels). 2-5 mice per group, bars show median with 95% CI. **(F)** Graphs depict potential patterns of autoreactive ASC recruitment: sporadic vs. continuous (top), and potential longevity patterns of the autoantibody response: short-lived vs. long-lived cohorts (bottom), based on Flag^+^ and Strep^+^ serum autoantibody titres, respectively. **(G-H)** Flag^+^ and Strep^+^ anti-DNA antibody titres in *Tlr7*^GoF^, (G) *Trex1^- /-^*, and *Dnase1l3^-/-^* (H) *Cd79a-Igk^Tag^* mice after tamoxifen. Median of 9 mice (G) and 13 and 9 mice respectively (H). Individual values are shown in Supplemental Figure 4B. For *Trex1^-/-^ Cd79a-Igk^Tag^* mice, time points have been grouped into 2-week intervals. **(I)** anti-DNA Flag^+^ and Strep^+^ antibody titres in 5-week-old and 12-week-old mice at the time of tamoxifen treatment. Median of 3-5 mice shown, representative of 3 experiments; individual values shown in Supplemental Figure 4F. **(J)** The recruitment index of Flag^+^ divided by total anti-DNA antibody titres (Flag^+^ / (Flag^+^ + Strep^+^) calculated at consecutive timepoints after tamoxifen in 5-week- and 12-week-old *Cd79a-Igk^Tag^ Tlr7*^GoF^ mice. 3 *Tlr7*^GoF^ mice per group shown, representative of 3 experiments. **(K)** Flag^+^ and Strep^+^ anti-sm/RNP and anti-Ro60 antibody titres in *Tlr7*^GoF^ *Cd79a-Igk^Tag^* mice after tamoxifen. Lines show median of 9 mice, respectively; individual values are shown in Supplemental Figure S4I. **(L)** Flag^+^ and Strep^+^ total antibody titres in *Tlr7*^GoF^ *Cd79a-Igk^Tag^* and *Tlr*^WT^ *Cd79a-Igk^Tag^* mice. Thin lines represent individual values, broad lines represent median.

Since IgG autoantibodies are considered the most pathogenic in SLE (Peng, Szabo, and Glimcher 2002), we first confirmed that isotype switching does not interfere with detection of the Igκ-tag (Figure S1C-H). When gated on total splenic B cells, Igκ-tag detection in IgG1^+^ and IgG2c^+^ cells remained close to 50% in *Igk^TagWT+^* mice, corresponding to full (∼100%) detection of *Igk^Tag^*-expressing cells (Figure S1C, F). The detection efficiency of the Igκ-tag was slightly lower in germinal centre B cells (∼ 80% of *Igk^Tag^* cells; Figure S1 D, G) and plasma cells (∼60% of *Igk^Tag^*cells; Figure S1E, H). Within each subset, the proportion of Flag-tagged cells was comparable across IgA-, IgM-, and IgG1-expressing populations (Figure S1C–H).

To investigate the longevity and replenishment of self-reactive B cells in lupus, we took advantage of a recently generated lupus mouse model driven by a *Tlr7* gain-of-function (GoF) genetic variant found in a child with SLE (Brown et al. 2022). As previously shown (Brown et al. 2022), *Tlr7*^GoF^ mice displayed increased frequencies of plasma cells in both spleen and bone marrow, although total plasma cell numbers in the bone marrow were comparable to *Tlr7*^WT^ controls (Figure S2A-B, top panels). In WT mice, approximately 50% of bone marrow plasma cells lacked CD19 and B220 expression, whereas this double-negative fraction was higher (∼60%) in *Tlr7*^GoF^ mice (Figure S2B, bottom panel), with a similar trend observed in the spleen (Figure S2A, bottom panel). To ensure that the *Igk*^Tag^ allele did not alter the autoimmune phenotype, we compared spleen weight, frequencies of CD11c^+^ B cells and plasma cell, and anti-DNA antibody titres among *Tlr7*^GoF^ mice with *Igk^WT/WT^*, *Igk^WT/Tag^* or *Igk^Tag/Tag^* genotypes. None of these parameters were affected by the presence of the *Igk^Tag^* allele (Figure S2C-E).

To track the turnover of B cell populations, we crossed *Tlr7*^GoF^ mice with *Cd79a-Igk^Tag^* mice and compared them to *Tlr7*^WT^ *Cd79a-Igk^Tag^* controls. Eight days after tamoxifen administration, *Igk^Tag^* splenic B cells had undergone tag-switching from Flag to Strep (Figure 1A-C and S3A). By day 30 and 52, the B cell pool showed gradual replenishment with newly generated Flag^+^ cells, which was nearly complete by day 52 in both *Tlr7*^WT^ and *Tlr7*^GoF^ mice (Figure 1C and S3A). Analysis of B cell subsets revealed higher Strep^+^ frequencies among CD11c^+^ age-associated B cells (ABCs) in *Tlr7^GoF^* mice (60% and 30% of Igκ^+^ cells at day 30 and 52, respectively) compared with *Tlr7*^WT^ mice (50% and 10%, respectively), suggesting slower ABC turnover in *Tlr7^GoF^*mice (Figure 1D). Conversely, the turnover of bone marrow plasma cells appeared faster in *Tlr7*^GoF^ mice, with 30% Strep^+^ plasma cells at day 52 compared to 50% in *Tlr7*^WT^ mice (Figure 1E).

In a complementary approach, we generated *Tlr7*^GoF^ and *Tlr7^WT^* Cd79^ERT2cre^ mice that express the fluorescent reporter mT/mG (Muzumdar et al. 2007). In these mice, Cre-mediated excision of tdTomato allows expression of a membrane-targeted GFP. Consistent with Igκ-tag results, by day 50 after tamoxifen administration, *Tlr7*^GoF^ mice showed higher frequencies of GFP^+^ (Strep^+^ equivalent) ABCs compared to *Tlr7*^WT^ control mice, whereas GFP^+^ plasma cells in spleen and bone marrow were reduced in *Tlr7*^GoF^ mice (Figure S3B-G).

### Continuous recruitment of newly-produced self-reactive B cells

Emergence of Flag-tagged anti-DNA antibodies after tamoxifen-induced Flag-to Strep-tag switching indicates recruitment of newly generated self-reactive B cells from the bone marrow into the autoantibody-secreting pool. Using this principle, we can follow recruitment patterns of Flag^+^ autoreactive cells and determine whether recruitment occurs sporadically (e.g., upon insult) or continuously (Figure 1F).

We labelled B cells in *Tlr7*^GoF^ *Cd79a-Igk^Tag^*mice by tamoxifen administration (inducing Flag-to Strep-tag switching) and followed the anti-DNA antibody response for 215 days by regular serum collection (Figure 1B). Whereas Cre^-^ or untreated *Tlr7*^GoF^ *Cd79a-Igk^Tag^* mice did not develop detectable Strep^+^ anti-DNA antibodies (Figure S4A), tamoxifen administration efficiently fate-mapped the majority of the autoantibody secreting cells in *Tlr7*^GoF^ *Cd79a-Igk^Tag^* mice reflected in ∼80% Strep^+^ anti-DNA circulating antibodies at early time points after tamoxifen (Figure 1G and S4B-C). *Cd79a-Igk^Tag^* mice expressing WT TLR7 showed negligible levels of anti-DNA antibodies of either tag (Figure S4B).

To capture the reciprocal relationship between decreasing Strep^+^ titres and increasing Flag^+^ titres, we calculated a “recruitment index” by dividing Flag^+^ by total Flag^+^ and Strep^+^ titres *(Flag/(Flag+Strep))*. By day 200 post-fate mapping, the median recruitment index for anti-DNA antibodies reached approximately 50%, demonstrating that over half of the autoantibody-secreting cell pool had been replaced by newly recruited cells (Figure 1G and Figure S4C). To evaluate whether these insights hold true more generally, we analysed serum anti-DNA titres in two additional mouse models carrying loss-of-function variants in *TREX1* and *DNASE1L3*, both established genetic causes of human SLE (Vinuesa, Shen, and Ware 2023). We observed similar overall kinetics of Strep-vs. Flag-labelled autoantibodies, though *Trex1*^-/-^ and *Dnase1l3^-/-^* mice showed a tendency towards slower turnover of the autoantibody response, reflected by lower recruitment indices at most time points compared with *Tlr7*^GoF^ mice (Figure 1G-H and S4B-D).

Anti-DNA autoantibody titres in *Tlr7*^GoF^ mice fluctuate significantly throughout the lifetime of the mouse and between individual mice, as do Flag- and Strep-tagged anti-DNA antibodies in tamoxifen-treated *Cd79a-Igk^Tag^ Tlr7*^GoF^ mice (Figure S4B and S4E). To evaluate whether age explains some of this variability, we stratified mice by age at the time of tamoxifen administration.

Tamoxifen administration to 5-week-old mice led to early emergence of Flag^+^ anti-DNA antibodies (produced by newly recruited B cells), which accounted for approximately 70% of the anti-DNA response by week 12. In older mice (12 weeks old at time of tamoxifen treatment), Flag^+^ antibodies were detected only after 8 weeks and accounted for approximately 25% of the response by week 12. We calculated the recruitment index for both age groups in *Tlr7*^GoF^ and *Trex1^-/-^* lupus-prone mouse models, which confirmed more efficient turnover of self-reactive B cells in young mice across both models, compared with older mice (Figure 1I-J and Figure S4F-G). To determine the recruitment of Flag^+^ cells independently of the pre-existing response (Strep^+^) we also calculated the relative increase in Flag^+^ antibodies after tamoxifen-induced tag-switching. Young mice of both *Tlr7*^GoF^ and *Trex1^-/-^* models showed higher titres and earlier appearance of Flag^+^ antibodies compared to older mice, indicating that the recruitment of newly generated cells into the anti-DNA antibody response is more efficient in young mice (Figure S4H).

### Sustained production of autoantibodies by a long-lived B cell cohort

We next examined the longevity of the self-reactive B cell cohort by measuring the persistence of Strep^+^ autoantibodies derived from B cells that were fate-mapped at the time of tamoxifen administration (Figure 1F). Remarkably, and despite the loss of Strep-labelled plasma cells over time in *Tlr7*^GoF^ mice (Figure 1E and S3E-G), Strep^+^ anti-DNA antibody titres were maintained for over 200 days with only a small decline (Figure 1G), suggesting persistence of a Strep-labelled cell cohort that sustains autoantibody production. Autoantibody responses against RNA-associated proteins, sm/RNP and Ro60, were also long-lived. However, they appeared to recruit new self-reactive cells more rapidly than the anti-DNA response. This was reflected by stable Strep^+^ autoantibody titres after fate-mapping and a faster increase in Flag^+^ antibodies, reaching 50% of the response by day 100, compared to day 200 for anti-DNA antibodies (Figure 1K and Figure S4I).

To examine how the kinetics of the autoantibody response compare to the total (or natural) antibody pool, we analysed total Flag^+^ and Strep^+^ titres in *Tlr7*^GoF^ and control *Tlr7*^WT^ mice. *Tlr7*^GoF^ mice had overall higher antibody titres than *Tlr7*^WT^ mice. Similar to the anti-DNA, anti-sm/RNP and anti-Ro60 autoantibody response, total Strep^+^ immunoglobulin titres were sustained through day 200 post fate-mapping regardless of *Tlr7* genotype (Figure 1L). Flag^+^ antibody titres increased steadily after fate-mapping in both *Tlr7*^WT^ and *Tlr7*^GoF^ mice and recruitment indices were comparable over time, independent of genotype or age at the time of tamoxifen administration (Figure 1L and S4J-K). Sustained Strep^+^ titres together with increasing Flag^+^ titres reflect the overall increase in total Ig as mice age.

Together, our data suggest that the recruitment of self-reactive B cells is an ongoing process and that B cell cohorts (Strep^+^ labelled) may be able to sustain the autoantibody response over extended periods of time.

### The autoantibody response is not sustained by long-lived plasma cells [Figure 2]

A sustained autoantibody response from a time-stamped B cell cohort (Figure 1G) can stem from either: (1) a population of long-lived plasma cells, typically thought to reside in the bone marrow, or (2) a proliferating pool of autoreactive B cells that continuously replenishes autoantibody-secreting cells. To distinguish between these possibilities, we fate-mapped plasma cells in *Igk^Tag^ Tlr7*^GoF^ mice using two different inducible Cre recombinases – *Prdm1^ERT2cre^* and *Jchain^ERT2cre^* – that specifically label antibody-secreting cells, and followed autoantibody secretion from these labelled populations over time (Figure 2A).

**Figure 2:**
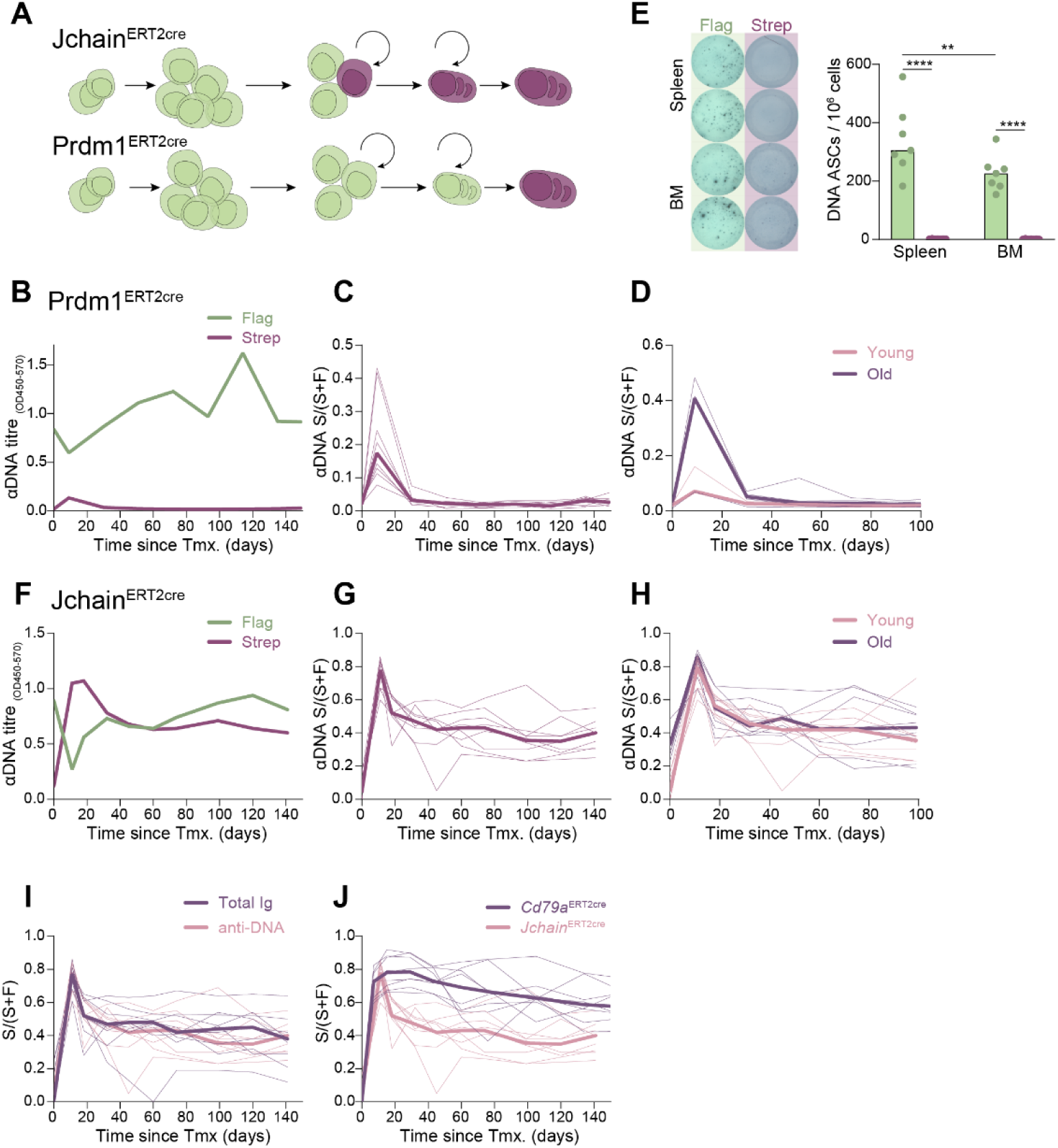
The autoantibody response is not sustained by long-lived plasma cells. **(A)** Schematic illustration of Cre-mediated tag switching of antibody-secreting cells from Flag^+^ (green) to Strep^+^ (purple) upon tamoxifen administration via *Prdm1^ERT2cre^* and *Jchain^ERT2cre^*. **(B-D)** Anti-DNA Flag^+^ and Strep^+^ antibody titres (B) and retention index (Strep^+^ / [Flag^+^ + Strep^+^]) (C) in *Tlr7*^GoF^ *Prdm1^ERT2cre^ Igk^Tag^* mice post-tamoxifen. (B) Median value of 8 mice; individual values in Supplemental Figure S5A. (C) Retention index for all mice. (D) Retention index stratified by age: young (4-5 weeks) vs. older (9 weeks). 3-4 mice per group; representative of two experiments. **(E)** ELISPOT quantification of Flag^+^ and Strep^+^ anti-DNA antibody-secreting cells in spleen and bone marrow at 100 days post-tamoxifen in *Tlr7*^GoF^ *Prdm1^ERT2cre^ Igk^Tag^* mice. 7 mice analysed; bars indicate median. **(F-H)** Flag^+^ and Strep^+^ anti-DNA antibody titres (F) and retention index (G-H) in *Tlr7*^GoF^ *Jchain^ERT2cre^ Igk^Tag^* mice post-tamoxifen. (F) Median of 9 mice; individual values in Supplemental Figure S5A. (G) Retention index for all mice. (H) Retention index stratified by age: young (5-6 weeks) vs. older (8-13 weeks), 8 mice per group across two experiments. **(I)** Retention index of total (Igĸ^+^) versus anti-DNA antibodies in *Tlr7*^GoF^ *Jchain^ERT2cre^ Igk^Tag^* mice. **(J)** Comparison of retention indeces post-tamoxifen in *Tlr7*^GoF^ *Cd79a-Igk^Tag^* versus *Tlr7*^GoF^ *Jchain-Igk^Tag^* mice. Thin lines represent individual values; thick lines represent medians.

*Prdm1^ERT2cre^* is active in cells that express the transcription factor BLIMP1 which drives B cell differentiation to plasma cells (Nutt et al. 2015). J-chain is an IgA/IgM multimerization factor, which is reliably transcribed in all plasmablast and plasma cell subsets from the pre-plasmablast stage. *Jchain^ERT2cre^*-mediated genetic manipulation is limited to plasmablasts and plasma cells (Xu, Barbosa, and Calado 2020) (Figure 2A).

Surprisingly, tamoxifen administration to *Tlr7*^GoF^ *Prdm1 ^ERT2cre^ Igk^Tag^* mice induced Flag-to-Strep switching in only 30-40 % of anti-DNA antibody-secreting cells in 9-week-old mice and 10-20 % in 4-5-week-old mice by day 7 (Figure 2B-C and S5A). Because plasmablasts express lower levels of BLIMP1 than plasma cells (Kallies et al. 2004), this suggests that a significant fraction of autoreactive antibody-secreting cells are plasmablasts lacking Prdm1 expression. Alternatively, autoreactive plasma cells may transcribe less *Prdm1* than those formed against exogenous antigens, resulting in reduced Cre-mediated recombination efficiency.

Strikingly, no Strep^+^ anti-DNA antibodies were detected 3-4 weeks after tamoxifen-induced Flag-to-Strep switching, suggesting that all anti-DNA antibody-secreting plasma cells are short-lived. Indeed, by day 30 onwards, the anti-DNA response consisted entirely of Flag^+^ antibodies derived from newly formed antibody-secreting cells or from cells resistant to tamoxifen-induced recombination (Figure 2B-D and S5A). Consistent with this, ELISPOT quantification of DNA-specific antibody-secreting cells in spleen and bone marrow confirmed the absence of long-lived plasma cells, as Strep^+^ cells were undetectable 100 days after tamoxifen administration (Figure 2E).

To fate-map all plasmablasts and plasma cells, including plasma cells with low *Prdm1* transcription, we used the *Jchain^ERT2cre^*model. Unlike Prdm1-driven labelling, *Jchain^ERT2cre^* fate-mapped ∼90% of autoantibody secreting cells (Figure 2F-H and S5A). Yet, similar to the *Prdm1-Igk^Tag^* model, Strep^+^ anti-DNA titres declined rapidly, with only 50-60% remaining by day 20 (Figure 2G). The Strep^+^ fraction declined at comparable rates in young mice and adult mice (Figure 2H) and in anti-DNA and total Ig responses (Figure 2I), indicating that plasmablast turnover is unaffected by age or antigen-specificity.

To evaluate the contribution of Jchain-expressing cells to the persistent Strep^+^ autoantibody response seen in *Cd79a-Igk^Tag^* mice, we compared retention index curves from *Jchain-Igk^Tag^*and *Cd79a-Igk^Tag^* models (Figure 2J). This analysis revealed that only about half of the sustained anti-DNA response was captured by Jchain^ERT2cre^ labelling (Figure 2J). Consequently, while plasmablasts and plasma cells contribute to autoreactive antibody production, a significant proportion of the cells sustaining the response must be Jchain-negative. Taken together, our results suggest that autoreactive plasma cells in the *Tlr7*^GoF^ model are short lived, and that the autoantibody response is sustained by a combination of Jchain-positive B cells – i.e., plasmablasts – and Jchain-negative precursors, such as ABCs, that continuously replenish the plasmablast and plasma cell pools.

### A population of (spleen-resident) proliferating cells express both ABC and PB markers [Figure 3]

To investigate the proliferative activity and properties of B cell subsets likely contributing to the self-reactive ASC pool, we performed single-cell RNA sequencing of ABCs and plasmablasts/plasma cells from spleen and bone marrow. To distinguish long-lived cohorts from newly recruited cells, we used *Tlr7*^GoF^ *Cd79^ERT2cre^ mT/mG* mice, in which the *mT/mG* allele enables fate-mapping based on fluorescence. We treated three *Cd79a^ERT2cre^ -mTmG Tlr7*^GoF^ mice with tamoxifen, and 90 days later, sorted GFP^+^ (Strep^+^ equivalent) and Tomato^+^ (Flag^+^ equivalent) splenic plasma cells (i), bone marrow plasma cells (ii) and splenic CD11c^+^ B cells (ABCs) (iii) into five 384-well plates (Figure 3A). Single-cell RNA sequencing was performed using an adapted FLASH-seq protocol (Hahaut et al. 2022) on a total of 1910 cells. After confirming uniform clustering per mouse, the data were pooled and analysed as a single dataset. Umap-clustering identified eleven distinct clusters which were annotated based on isotype expression and cell type (Figure 3B-E and S5B).

**Figure 3:**
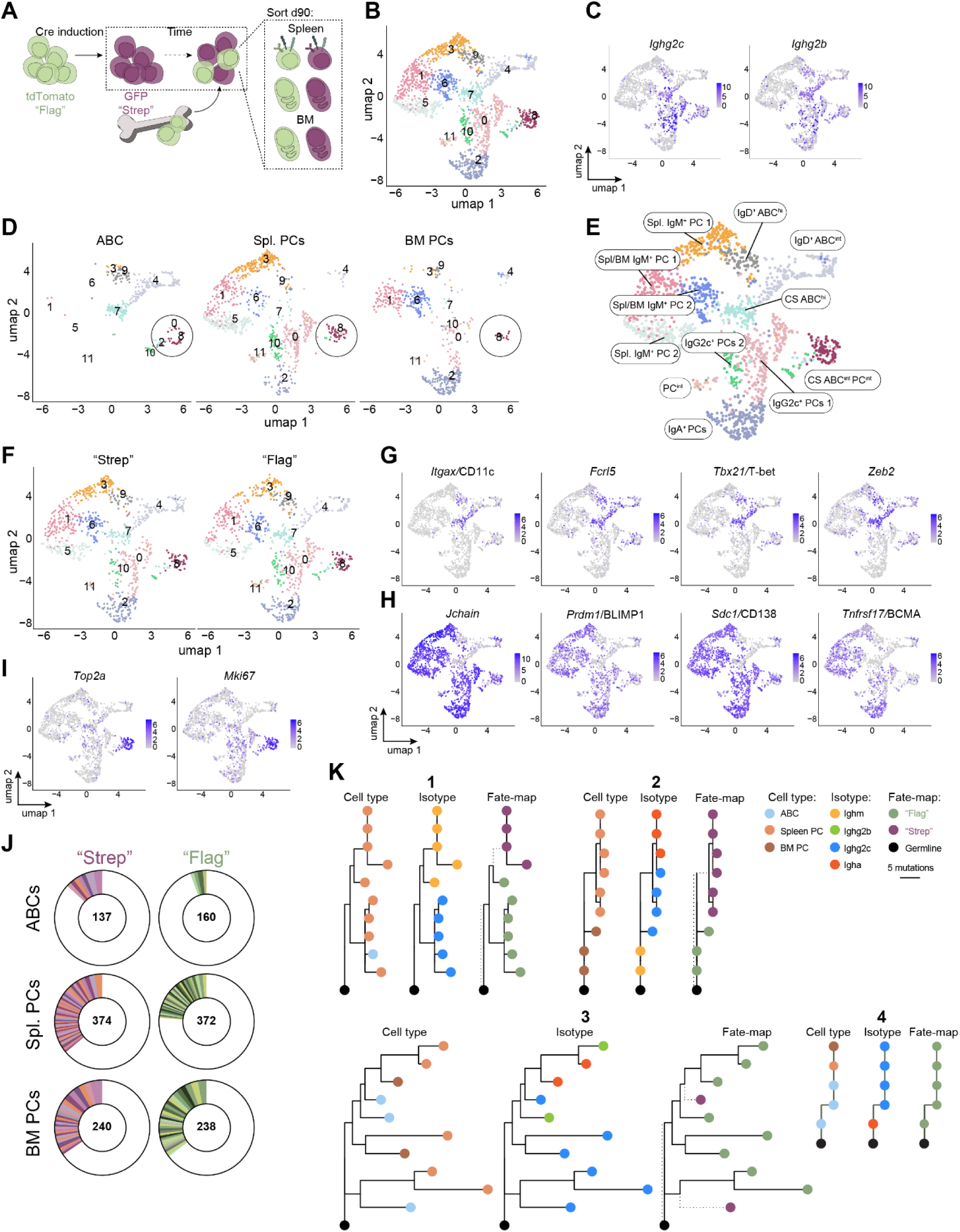
ABCs and plasmablasts are transcriptionally and clonally related. **(A)** Schematic showing the experimental setup and illustration of the cell types sorted at day 90 post-tamoxifen. From each cell type both tdTomato^+^/ “Flag” (green) and GFP^+^/ “Strep” (purple) cells were sorted. A total of 1910 cells were sequenced and 1826 cells passed the quality control criteria: 190 GFP^+^ ABCs, 187 tdTomato^+^ ABCs, 265 GFP^+^ bone marrow (BM) PCs, 270 tdTomato^+^ BM PCs, 464 GFP^+^ spleen PCs, 450 tdTomato^+^ PCs **(B)** Umap and clustering analysis of all ABCs and plasmablasts from spleen and bone marrow. **(C)** Expression of immunoglobulin isotype Ighg2c and Ighg2b transcripts. **(D)** umap from C split by sorting niche. Circles mark cluster 8. **(E)** Umap annotated by sorting niche and isotype switching status. **(F)** Umap from C split by early (GFP^+^ / “Strep”) and late (Tomato^+^ / “Flag”) recruitment. **(G-I)** Expression of ABC markers (G), plasmablast markers (H), and proliferation markers (I). **(J)** Clonality within GFP^+^ / “Strep” and tdTomato^+^ / “Flag” cells from each sorting niche. White = single cell clones, coloured = 2+ cells per clone, increase in size clockwise. Number in the centre of the circle plot = number of cells analysed. **(K)** Clonal trees based on H-chain similarity within single clonal families. Each clonal tree is split into three and coloured by: Cell type (left), Isotype (centre), and tdTomato / “Flag” or GFP / “Strep”. The length of horizontal lines represents the mutational distance.

When stratified by GFP (Strep^+^ equivalent) or tdTomato (Flag^+^ equivalent), cluster composition showed minimal differences (Figure 3F). This suggests that the length of time elapsed since ABC or plasma cell recruitment in lupus-prone *Tlr7*^GoF^ mice does not influence their differentiation state and by inference, their ability to feed into the population giving rise to autoantibody-secreting plasma cells. CD11c^+^ B cells were distributed across four clusters, all of which expressed ABC markers (*Itgax*, *Tbx21*, *Fcrl5*, *Zeb2)* (Figure 3D and 3G) and contained both class-switched and IgM/IgD-expressing cells (Supplemental Figure S5C). Most plasma cell clusters were shared between spleen and bone marrow, but their relative abundance differed across tissues. Consistent with prior reports, IgA^+^ plasma cells were primarily bone marrow-resident, whereas IgG2c class-switched plasma cells — associated with T-bet- and IFN-g-driven autoimmunity (Stone et al. 2019; Dai et al. 2024) — were most abundant in the spleen (Figure 3C-E and S5B).

Intriguingly, one cluster (cluster 8) was present across all three sorted populations (i-iii above) and contained IgG2b and IgG2c class-switched cells along with markers of ABCs and plasma cells (Figure 3C-D and 3G-H). Cell-cycle gene analysis revealed up to six-fold higher expression of S/G2/M-phase genes *Mki67* and *Top2a* compared with other clusters (Figure 3I). Interestingly, at the cluster periphery, cells expressed ABC markers with low PC marker levels, whereas cells at the centre resembled *bona fide* plasmablasts. This pattern may indicate that some ABCs within the cluster may be differentiating toward a plasmablast fate, consistent with previous reports (Jenks et al. 2018). Notably, few cells in this cluster expressed *Tnfrsf17* (encoding BCMA, a common plasma cell marker and CAR-T target) (Figure 3G-H), and *Cd19* expression was low in both splenic and bone marrow plasma cells (Supplemental Figure 5D).

Together, these results identify a spleen-resident population of class-switched, highly proliferative ABCs and plasmablasts that cluster closely together. We hypothesize that these *Jchain*^-^ ABCs and Jchain^+^ plasmablasts (Figure 3H), along with counterparts in other tissues, act as a precursor pool that continuously replenishes autoantibody-secreting cells over time. To explore the relatedness between splenic ABCs and plasma cells in spleen and bone marrow, we analysed their BCR repertoires. We first assessed clonal expansion within the GFP^+^ (Strep^+^ equivalent, induced by tamoxifen administration) and tdTomato^+^ (Flag^+^ equivalent, newly recruited) fractions. Both splenic ABCs and plasma cells showed greater clonality in the GFP^+^ fraction (Figure 3J), suggesting that although early- and late-recruited cells are transcriptomically similar (Figure 3F), certain VDJ combinations may confer a selective survival advantage. In contrast to splenic plasma cells, bone marrow plasma cells did not exhibit differences in clonality based on time of recruitment (Figure 3J).

To identify clonal relationships between ABCs and plasma cells, we examined four of the largest expanded clones that contained IgG2c class-switched cells. These clones differed in the extent of somatic hypermutation (SHM) and clonal diversity: Clone 4 consisted of one unmutated cell and four cells with two mutations within CDR2, whereas Clone 3 had up to 13 replacement mutations including some within CDR3 (Figure 3K). The median replacement-to-total mutation ratio was 0.54 suggesting weak selection pressure acting on somatic hypermutation. Strikingly, we observed clonal overlap between cells sorted as ABCs and splenic and bone marrow plasma cells, suggesting that ABCs give rise to plasma cells in both locations (Figure 3K). In addition to IgG2c^+^ cells, IgA, IgM and IgG2b class-switched cells were also present within individual clones, demonstrating that autoantibodies of different isotypes can originate from the same common ancestor. Finally, three of the four clones contained both GFP^+^ (Strep^+^ equivalent) and tdTomato^+^ (Flag^+^ equivalent) cells. Given that tamoxifen-induced labelling efficiency was 80-85%, these clones likely arose independently on at least two occasions, suggesting preferential selection of certain H-L chain combinations into the IgG2c^+^ ABC and plasma cell repertoires in *Tlr7*^GoF^ mice (Figure 3K).

We conclude that ABCs differentiate into plasma cells of different isotypes that migrate to various locations, including the spleen and the bone marrow. Furthermore, despite an increase in overall clonality over time, there appears to be limited selective pressure on individual clones.

### The spleen serves as a reservoir of autoreactive ASCs [Figure 4]

While bone marrow plasma cells are thought to be the primary source of circulating autoantibodies, secondary lymphoid organs and inflamed tissues including the kidneys have been suggested to be enriched in autoreactive plasma cells in some lupus mouse models (Steinmetz et al. 2023; Starke et al. 2011; Hoyer et al. 2004; Mathian et al. 2011).

We therefore next sought to define the tissues supporting autoreactive ASCs across three mouse models of human SLE. Given that mucosal-derived IgA anti-DNA antibodies may interact with IgG to mediate effector functions (Waterman et al. 2024) and that salivary glands and draining submandibular lymph nodes are enlarged in *Tlr7*^GoF^ mice, we included these lymph nodes as well as Peyer’s Patches in our analysis. Quantification of self-reactive IgG ASCs by ELISPOT in WT, *Tlr7*^GoF^, *Dnase1L3*^-/-^, and *Trex1*^-/-^ mice revealed that the spleen contained the highest frequency of anti-DNA-secreting cells across all models (out of 10^6^ cells) (Figure 4A-B). Bone marrow contained the second highest frequency, with 58%, 72%, and 12% fewer anti-DNA ASCs than spleen in *Tlr7*^GoF^, *Dnase1l3*^-/-^ and *Trex1^-/-^*mice, respectively.

**Figure 4:**
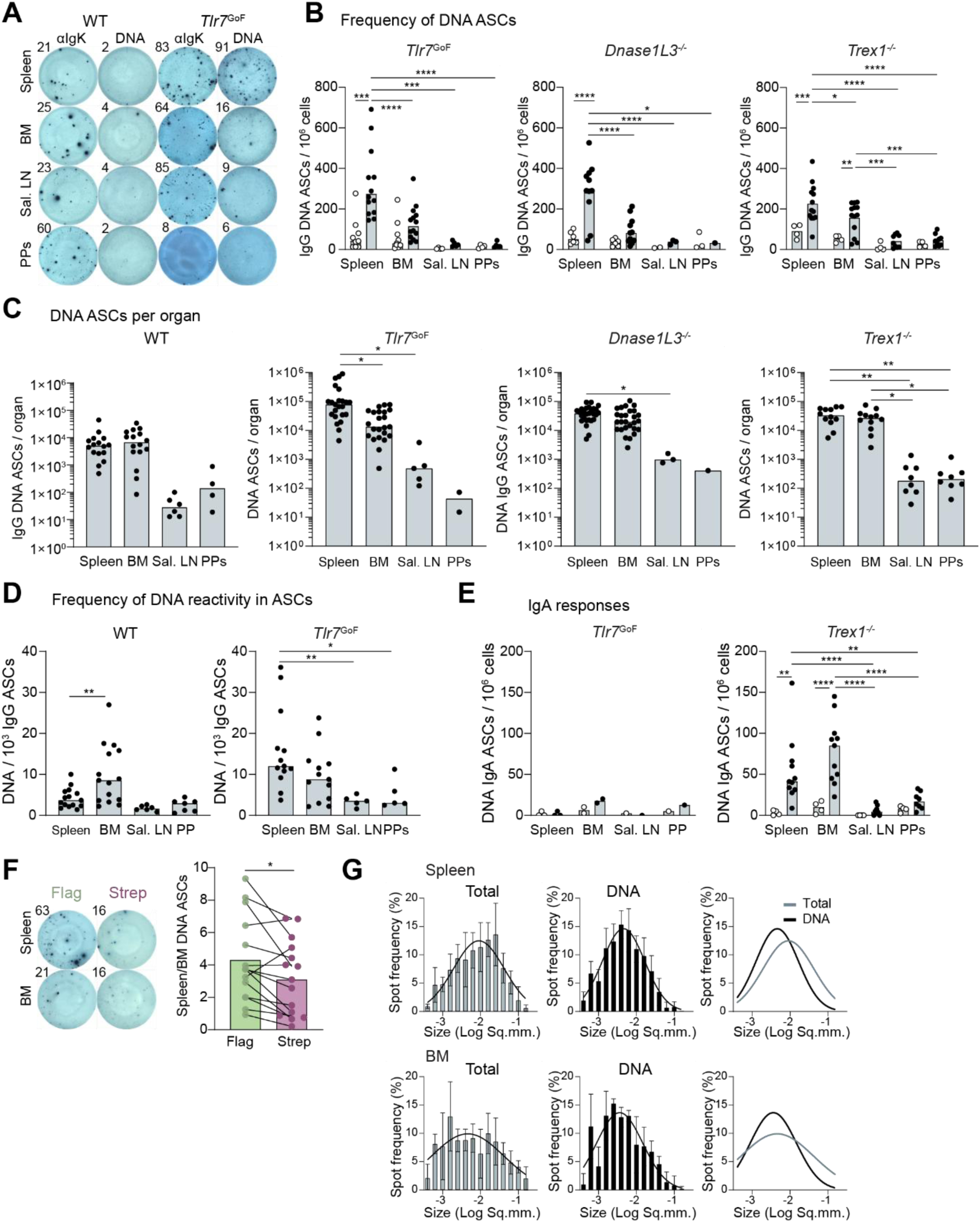
The spleen is the primary site for enrichment of autoreactive ASCs in lupus models. **(A)** Representative ELISPOT for total (Igĸ^+^) and DNA-reactive antibody-secreting cells (ASCs) in tissues isolated from WT and *Tlr7*^GoF^ mice. **(B)** Quantification of IgG^+^ DNA-reactive ASCs as a fraction of total cellularity from ELISPOT as in A. Three different models of monogenic lupus (black dots) analysed in parallel with WT control mice (empty dots). Pooled data from 3-4 assays, representative of 2+ pooled experiments. **(C)** Estimated numbers of IgG^+^ DNA-reactive ASCs per organ for three models of monogenic lupus. **(D)** Frequency of DNA-reactivity in the ASC population in four different tissues in WT and Tlr7^GoF^ mice. **(E)** Quantification of IgA^+^ DNA-reactive ASCs as a fraction of total cellularity in *Tlr7*^GoF^ and *Trex1^-/-^* mice in parallel with WT mice, from ELISPOT assays as in A. Data pooled from 2-3 experiments. **(F)** ELISPOT of Flag^+^ and Strep^+^ DNA-reactive ASCs in the spleen and bone marrow of *Tlr7*^GoF^ mice. Quantification shows relative proportion of cells in spleen vs bone marrow (right). **(G)** ELISPOT Spot size quantification of total (Igĸ^+^) and DNA-reactive ASCs in spleen (top) and bone marrow (bottom). Pooled data from 2 assays, representative of 2+ pooled experiments. PPs = Peyer’s Patches, Sal. LN = Salivary lymph nodes, BM = Bone marrow.

To estimate the total number of ASCs per organ, we calculated bone marrow cellularity assuming that one femur and tibia together represent 9.7% of total bone marrow cells (Mahajan et al. 2015). Using this approach, only *Tlr7*^GoF^ mice exhibited an increase in total DNA-ASCs cells in the spleen relative to bone marrow (5.9-fold, Figure 4C). *Tlr7*^GoF^ mice also trended toward higher numbers of IgG DNA-ASCs in submandibular lymph nodes compared with WT mice (17-fold increase, p=0.09) (Figure 4C). When expressed as a fraction of total IgG ASCs, anti-DNA ASCs were also most frequent in the spleen of *Tlr7*^GoF^ mice, whereas in WT mice, the highest frequency of DNA-reactive ASCs was observed in the bone marrow (Figure 4D).

In contrast to the splenic enrichment of IgG anti-DNA responses, IgA anti-DNA ASCs were predominantly found in the bone marrow, exhibiting an approximately 2-fold enrichment compared with the spleen (Figure 4E). This observation is consistent with the known enrichment of total IgA plasma cells in both mouse and human bone marrow (Wilmore et al. 2021). IgA anti-DNA ASCs were also detected in the Peyer’s patches and salivary lymph nodes of *Tlr7*^GoF^ mice, although their frequencies were not significantly higher than in WT mice (Figure 4E).

To examine whether DNA-ASCs generated early in life localise differently from newly cells, we induced Flag-to-Strep switching in all B cells via tamoxifen and performed ELISPOT assays on spleen and bone marrow over 100 days later. Both Strep^+^ cells (labelled by tamoxifen) and more recently recruited Flag^+^ cells preferentially localised to the spleen. However, the spleen-to-bone marrow ratio was higher for Flag^+^ DNA-ASCs than for Strep^+^ cells (Figure 4F), suggesting that over time, DNA-ASCs or their precursors either redistribute toward the bone marrow or have a survival advantage in this niche.

As a proxy for the quantity of antibody produced per cell, we measured the spot size of total (Igĸ^+^) and DNA-reactive ASCs. In both bone marrow and spleen, we found that the spot-size distribution of DNA-reactive ASCs was overall skewed towards smaller spots compared to total ASCs (Figure 4G). This suggests that DNA-reactive ASCs may produce lower quantities of antibody per cell compared with non-self-reactive ASCs and are thus less differentiated, which is consistent with our previous findings showing that anti-DNA ASCs are not long-lived plasma cells (Figure 3D-F). We conclude that the spleen is specifically enriched in autoreactive IgG ASCs and that these cells appear to exhibit a less terminally differentiated phenotype relative to the broader ASC population.

### ASCs in lupus-prone mice are frequently found surrounding B cell follicles [Figure 5]

After identifying the spleen as an important reservoir of self-reactive plasma cells, we next investigated their spatial distribution within this tissue by immunofluorescence. In WT mice, CD138^+^ plasma cells accumulated in bridging channels where white pulp meets the red pulp, and within the red pulp (Hargreaves et al. 2001; Toellner et al. 1996) (Figure 5A and S6A). In contrast, plasma cells in *Tlr7*^GoF^ mice were largely dispersed throughout the red pulp or formed clusters adjacent to CD19^+^ B cell follicles and marginal sinuses (the vascular space between the red pulp and white pulp where B cells sample antigen) (Figure 5B-E and S6B). Despite mouse-to-mouse variation in plasma cell localization, plasma cells were consistently absent from T cell zones (Fig. 5B-C).

**Figure 5:**
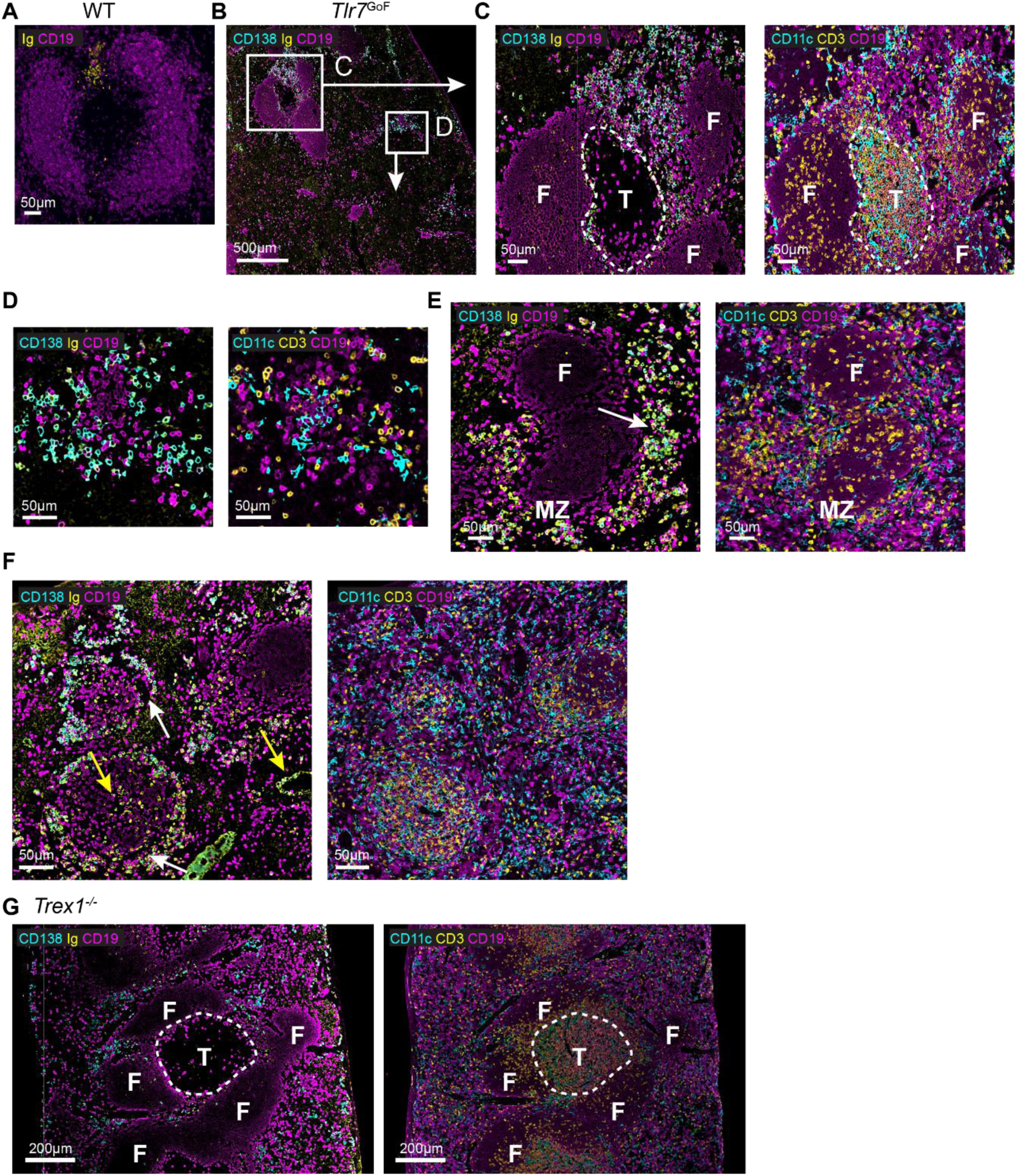
Plasma cell localization within the spleen of lupus-prone mice. **(A)** Representative immunofluorescence image of a WT spleen showing plasma cell localised in the bridging channel. **(B)** Representative image of a *Tlr7*^GoF^ spleen. White boxes indicate the areas shown in C and D. **(C-D)** Higher magnification views showing B cell follicles (“F”) and T cell zone (“T”) (C), and a plasma cell cluster within the red pulp (D). **(E)** Representative image of a *Tlr7*^GoF^ spleen showing a B cell follicle (“F”) and marginal zone (“MZ”); plasma cell clusters are indicated by a white arrow. **(F)** Representative image of circular structures in *Tlr7*^GoF^ spleen. Consecutive sections were stained for CD138, Ig, and CD19 (left) and for CD11c, CD3, and CD19 (right). White arrows indicate marginal zone sinuses; yellow arrows indicate vascular structures. **(G)** Representative images of *Trex1^-/-^* spleens showing a T cell zone (“T”) and B cell follicles (“F”). Left and right panels show staining for CD138, Ig, CD19 (left) and CD11c, CD138, CD19 (right) on consecutive sections.

Occasionally, we observed unique circular structures composed of interspersed B cells, T cells and dendritic cells, encircled by a ring of B cells and plasma cells, with the latter separated by a region resembling the marginal sinus (Figure 5F, white arrows). Plasma cells were also occasionally found in proximity to central arterioles or lined structures consistent with blood vessels (Figure 5F, orange arrows). Within the red pulp, plasma cells frequently localised near B cells, often in the absence of nearby dendritic cells (Figure 5B-D and Supplemental Figure S6B-E).

Further analysis of CD19, CD138, and Ig expression revealed that plasma cell clusters contained cells heterogenous populations, including cells expressing CD19 alone, CD19 together with Ig, triple positive CD19^+^ Ig^+^ CD138^lo^ cells, and CD138^+^ Ig^+^ double positive cells. This pattern indicates that cells along the plasma cell differentiation path — from CD19^+^ B cells to CD138^-^ CD19^+^ Ig^+^ plasmablasts and CD138^+^ Ig^+^ CD19^-^ plasma cells — exist within the same cluster and potentially mature in situ (Supplemental Figure S6F).

To broaden our findings to another model of monogenic lupus, we analysed spleens from *Trex1^-/-^* mice. Plasma cell localization in these mice resembled that of *Tlr7*^GoF^ mice, characterised by exclusion from T cell zones and by plasma cells dispersed throughout the red pulp, frequently clustering near B cell follicles, marginal zone sinuses, and vascular structures (Figure 5G).

In agreement with previous reports, Ig deposition was observed in the red pulp and within B cell follicles (marking follicular dendritic cells) of *Tlr7*^GoF^ mice (Figure S6B). Together, these results suggest that plasma cells localise uniquely in lupus-prone mice.

### Spleens of SLE patients contain abundant plasma cells [Figure 6]

We next investigated the presence and distribution of plasma cells in spleen samples from five patients with SLE, some of whom presented with well-recognized manifestations or overlapping conditions commonly associated with the disease, including immune thrombocytopenic purpura (ITP), autoimmune haemolytic anemia (AIHA), Sjögren’s syndrome, and antiphospholipid syndrome (APS) (Supplemental Table 1). One additional sample was obtained from a patient with isolated ITP. Five spleen samples resected for reasons unrelated to SLE were analysed as controls.

As previously described, spleens from SLE patients exhibited marked disruption of normal architecture, with poorly defined demarcation between T cell zones and B cell follicles, as well as an increased density of large vessels (Figure 6A-C) (N. Li et al. 2013). Plasma cells were more abundant in SLE spleens than in controls and, similar to mouse models, were dispersed throughout the red pulp. Notably, plasma cells frequently lined splenic blood vessels in both the white and red pulp, where they appeared to localise within the subendothelial space. In contrast, plasma cells in control spleens localised to T cell zones, bridging channels, and the red pulp, where they occasionally associated with blood vessels and capillaries (Figure 6A-C).

**Figure 6:**
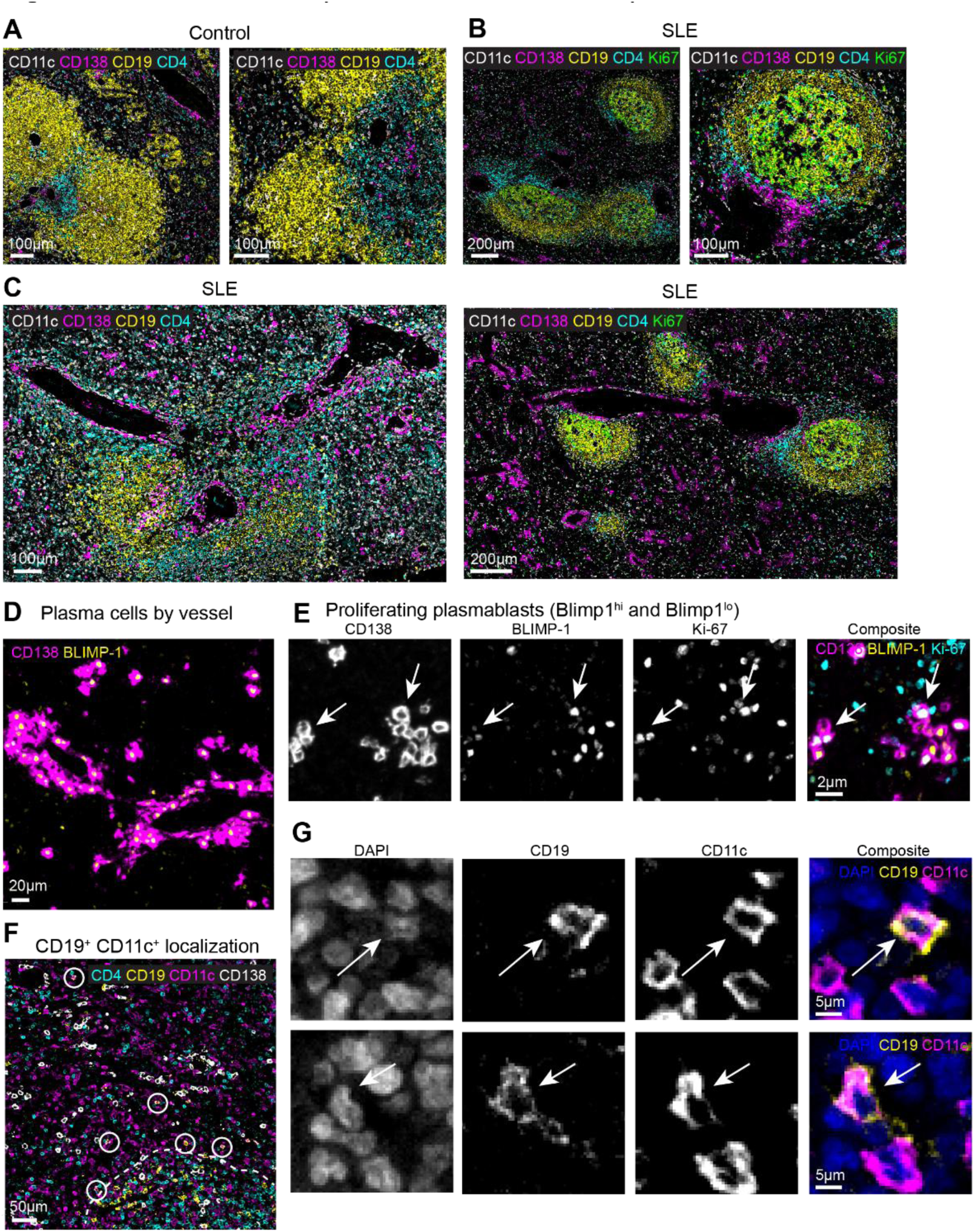
Plasma cell localization in spleens from SLE patients. **(A-C)** Immunofluorescence images of a representative spleens from a patient without autoimmune diagnosis (patient 4) (A) or from two patients diagnosed with SLE (patient 1 and 5) (B-C). **(D)** Plasma cells identified by CD138 and BLIMP1 staining around a vessel in the red pulp from a representative SLE spleen (Spleen 1). **(E)** Overlap between CD138, BLIMP1 and Ki67 staining in a cluster of plasma cells from an SLE spleen (Spleen 6). Arrows mark a BLIMP1^hi^ cell (right) and a BLIMP1^lo^ cell (left) respectively. Arrows mark a BLIMP1^hi^ cell (right) and a BLIMP1^lo^ cell (left) respectively. **(F)** Localization of CD11c^+^ CD19^+^ cells in a representative spleen from an SLE patient (patient 5). **(G)** Overlap between CD19 and CD11c staining from cells in F. Images are representative of five spleens from SLE patients and five spleens from control patients. Patient details are found in Supplemental Table 1.

We validated the plasma cell phenotype of CD138^+^ cells localised around blood vessels by confirming nuclear expression of BLIMP1 (encoded by *PRDM1*) in many of these cells (Figure 6D). Ki-67 staining further revealed that a subset of CD138^+^ cells (both BLIMP1^lo^ and BLIMP1^hi^) were actively cycling, consistent with a plasmablast phenotype (Figure 6E). As in mouse spleens, plasma cell density appeared higher in more vascularized regions towards the periphery of the spleen (Supplemental Figure 5G-I). In two of five spleens derived from SLE patients (patient 1 and 6), we observed germinal centre formation characterised by clusters of Ki67^+^ B cells within follicles (Figure 6B-C). In several instances, plasma cells accumulated around enlarged central arterioles separating the T cell zone from the dark zone of the germinal centre (a Ki-67^hi^, T cell depleted area). Limited numbers of plasma cells were found within the germinal centres themselves (Figure 6B). CD11c^+^ CD19^+^ cells were also observed in the red pulp and at higher frequencies around B cell / T cell enriched areas (Figure 6F-G). Actively cycling (Ki-67^+^) CD11c^+^ B cells were not readily visible in SLE spleens, suggesting either treatment-related suppression or an extra-splenic origin of plasma cells.

### CD19 CAR T cells are sufficient for full reset of lupus in Tlr7^GoF^ mice [Figure 7]

Our results so far indicate that the persistence of autoantibodies does not depend on long-lived plasmablasts or plasma cells, but rather on cycling ABCs or their precursors, residing in spleen and/or other tissues. This is supported by the observation that most anti-DNA ASCs do not persist beyond three weeks (Figure 3D-F). If this conclusion is correct, the following predictions should hold: i) Complete depletion of autoantibody-producing cells should be achievable by targeting ABCs and other CD19^+^ precursors. ii) Any re-emergence of autoantibodies after complete depletion of CD19^+^ B cells – e.g., in a relapse – should derive exclusively from B cells newly generated in the bone marrow, rather than from residual, non-depleted CD19^-^ B cells. iii) targeting BCMA, which is highly expressed on ASCs but not on ABCs (Martin et al. 2024), should not provide a therapeutic advantage to anti-CD19 treatment and may not achieve full immune reset. We then sought to test these predictions by depleting all CD19^+^ B cells from our lupus-prone mice with CAR-T cells.

To track immune reset, we used *Tlr7*^GoF^ *Cd79a-Igk^Tag^* mice and fate-mapped all B cells (Flag-to-Strep switch) via tamoxifen administration, followed by cyclophosphamide pre-conditioning and adoptive transfer of CD19-targeted CAR-T cells to achieve B cell depletion (Figure 7A). To investigate which cells drive relapse, we chose CAR-T cells expressing the murine 4-1BB co-stimulatory domain, which are known to be short-lived in mice (G. Li et al. 2018). Control mice received CAR-T cells specific for human IL13ra. As predicted (i), CD19-targeted CAR-T treatment led to complete depletion of circulating B cells by day 7, even in young mice with lower CAR^+^ T cell expansion (Figure 7C-E). Splenic B cells and plasma cells were also fully eliminated by week 4 (Figure 7F), likely as a result of both CD19 expression on a subset of plasma cells (Figure S2A-B), and their short lifespan (< 30 days) (Figure 2B-D). Correspondingly, autoantibody analysis showed complete loss of Flag^+^ and near-complete depletion of Strep^+^ anti-DNA antibodies within seven days of αCD19 CAR-T cell administration, indicating rapid antibody turnover (Figure 7G-I). A small, transient drop in both Flag^+^ and Strep^+^ autoantibody titres was detected at day 7 in control hIL13ra CAR-T treated mice, likely caused by the cyclophosphamide treatment (Figure 7G-I).

**Figure 7:**
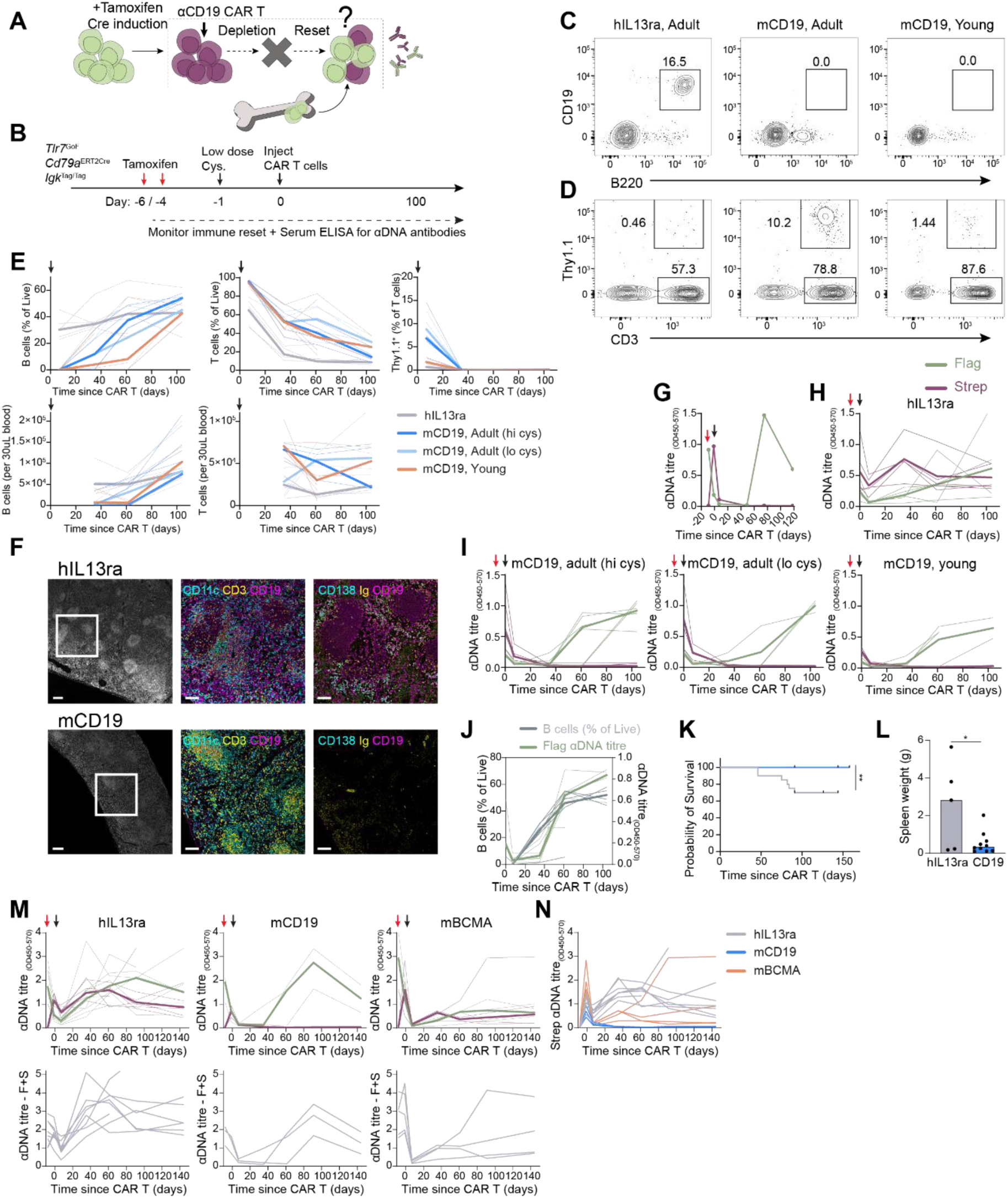
αCD19 CAR-T cells achieves complete immune reset in *Tlr7*^GoF^ mice. **(A)** Illustration indicating the rationale and potential outcome of CAR-T cell experiments. From left to right: Tamoxifen-induced tag-switching in *Tlr7*^GoF^ *Cd79a-Igk^Tag^* mice from Flag-(green) to Strep-tag (purple), B cell depletion by αCD19 CAR-T cells, and B cell reconstitution and recurrence of autoantibody-secreting cells expressing either Flag- or Strep-tag depending on origin. **(B)** Timeline of CAR-T cell experiments. **(C-D)** FACS analysis of peripheral blood 7 days after CAR-T cell transfer from adult mice (12-16w at time of CAR-T cell treatment) treated with αhuman IL13ra CAR-T cells (left), and αmouse CD19 CAR-T cells (middle), and young mice (5w at time of CAR-T treatment) treated with αmouse CD19 CAR-T cells (right). B and T cell staining shown in C and D, respectively. Thy1.1 indicates expression of the CAR. **(E)** Quantification of FACS analysis of B cells and T cells in peripheral blood as in C-D. Black arrows indicate time of CAR-T cell transfer. **(F)** Representative immunofluorescence images of *Tlr7*^GoF^ *Cd79a-Igk^Tag^* mouse spleens 35 days post treatment with human IL13ra-targeted (top) or mouse CD19-targeted (bottom) CAR-T cells. Left panels show DAPI staining of large spleen areas. White boxes indicate areas viewed in middle and right panels. Consecutive slides were stained for CD138, immunoglobulin (Ig), CD19 and CD11c, CD3, CD19. **(G)** Representative anti-DNA Flag^+^ and Strep^+^ antibody titres in a *Tlr7*^GoF^ *Cd79a-Igk^Tag^* mouse treated with tamoxifen (red arrow) and αCD19 CAR-T cells (black arrow). **(H-I)** anti-DNA Flag^+^ and Strep^+^ antibody titres in *Tlr7*^GoF^ *Cd79a-Igk^Tag^* mice treated with tamoxifen and CAR-T cells targeting human IL13ra (H) or mouse CD19 (I). (I) Adult mice (12-16 weeks at time of CAR-T transfer) treated with high dose (left) or low dose (middle) cyclophosphamide, and young mice (5 weeks at time of CAR-T transfer) treated with high dose cyclophosphamide (right). Red arrow = time of tamoxifen administration, black arrow = time of CAR-T transfer. **(J)** Simultaneous peripheral blood B cell percentage and anti-DNA Flag^+^ antibody titres after tamoxifen administration and αCD19 CAR-T cell transfer in *Tlr7*^GoF^ *Cd79a-Igk^Tag^* mice. **(K)** Survival data from 20 αhIL13ra-CAR-T treated and 17 αCD19-CAR-T treated *Tlr7*^GoF^ *Cd79a-Igk^Tag^* mice. **(L)** Spleen weight 140-150 days post CAR-T transfer in *Tlr7*^GoF^ *Cd79a-Igk^Tag^* mice treated with αhIL13ra- or αCD19-CAR-T cells. Representative of two experiments. **(M)** Top panels: Anti-DNA Flag^+^ and Strep^+^ antibody titres in *Tlr7*^GoF^ *Cd79a-Igk^Tag^* mice treated with tamoxifen and CAR-T cells targeting hIL13ra (left), mCD19 (middle) and mBCMA (right). Bottom panels: Sum of Flag^+^ and Strep^+^ anti-DNA titres. **(N)** Strep^+^ anti-DNA titres in individual *Tlr7*^GoF^ *Cd79a-Igk^Tag^* mice as in M. Thin lines represent individual mice, broad lines represent the median. “hi cys” = high cyclophosphamide dose (0.2mg/g), “lo cys” = low cyclophosphamide dose (0.1mg/g).

We next tested our second prediction — that autoantibodies reappearing after CAR-T cell treatment derive exclusively from newly generated bone marrow B cells. B cells gradually re-emerged between days 30 and 75 days post-treatment in adult mice and between days 40 and 80 days in young mice, coinciding with the decline of CAR^+^ T cells (Figure 7E). Anti-DNA antibodies were detectable within three weeks of B cell reconstitution in all mice, indicating that standard CAR-T cell treatment alone does not induce a durable immune reset in monogenic lupus (Figure 7J). Strikingly and as predicted, post-CAR-T cell anti-DNA antibodies were exclusively Flag^+^, confirming their origin from newly produced B cells, regardless of age or cyclophosphamide dose (Figure 7I). Furthermore, in four older mice that failed to reconstitute B cells by 140 days post-depletion, no autoantibodies of either tag were detected. This excludes the possibility that incomplete labelling of long-lived plasma cells in the *Cd79a-Igk^Tag^* system could sustain long-term autoantibody production (Supplemental Figure S7A).

*Tlr7*^GoF^ mice exhibit accelerated mortality (Brown et al. 2022) and although B cell reconstitution was seen from approximately 5 weeks after depletion, 100% of *Tlr7*^GoF^ mice treated with αCD19 CAR T cells survived up to 150 days after treatment compared to only 70% of *Tlr7*^GoF^ mice treated with hIL13ra CAR-T cells (Figure 7K). In addition, spleen weight at the endpoint (day 140-150 post-CAR-T) was higher in αhIL-13 treated control mice compared to αCD19 CAR-T treated mice (Figure 7L). These results demonstrate the efficiency of CD19-targeted CAR-T therapy in *Tlr7*-driven lupus, along with a high relapse rate in the presence of a strong genetic driver.

We then tested our third and final prediction: That there would be no advantage in treating mice with αBCMA CAR-T cells over αCD19 CAR-T cells, given the absence of long-lived plasma cells in *Tlr7*^GoF^ lupus and CD19 expression by ASC precursors. Three cohorts of age-matched mice were fate-mapped (Flag-to-Strep switch) by tamoxifen administration and treated with hIL13ra-, αCD19- or αBCMA-CAR-T cells in parallel. CAR^+^ T cells expanded comparably in αCD19- and αBCMA CAR-T treated mice 7 days after transfer, and anti-DNA antibody titres were depleted concomitantly in both mouse cohorts (Figure 7M and S7B).

Strikingly, while CD19-treated mice exclusively developed Flag^+^ autoantibodies after B cell reconstitution – indicating their origin from newly formed B cells – both Flag^+^ and Strep^+^ autoantibodies were detected in αBCMA CAR-T cell treated mice within 35 days of CAR-T cell transfer (Figure 7M-N). In line with our predictions, the reappearance of Strep^+^ autoantibodies shortly after BCMA-mediated plasma cell depletion suggests that αBCMA targeting therapy fails to deplete some autoreactive plasma cell precursors, which can replenish the pool of autoantibody-secreting cells.

We conclude that CD19-targeted CAR T cell therapy is sufficient to deplete all autoreactive antibody-secreting cells and their precursors across tissues. This is consistent with our finding that autoreactive antibody-secreting cells are short-lived rather than long-lived plasma cells and thus depletion of CD19-negative plasma cells is not required to achieve immune reset.

## Discussion

In this study, we combined a recently developed system to molecularly tag antibodies (Schiepers et al. 2023) with mouse models of lupus carrying human causal genetic variants to retrospectively trace the fate and behaviour of self-reactive cells producing tagged autoantibodies. This approach has allowed us to answer fundamental questions regarding the dynamics, longevity, and tissue localization of autoreactive B cell.

Despite identifying a long-lived cohort of autoreactive cells that continued to give rise to autoantibodies for over 200 days after a single tamoxifen injection, we found that the self-reactive plasma cells themselves appear short-lived. No BLIMP1^+^ plasma cell-derived autoantibodies were detectable within 30 days of labelling, the turnover of labelled plasma cells was shorter in *Tlr7*^GoF^ than in WT mice, and CAR-T cell mediated depletion of CD19^+^ cells efficiently depleted all plasma cells and autoantibodies within five weeks of CAR-T cell transfer. The rapid loss after CAR-T therapy likely reflects the combined effects of eliminating CD19^+^ plasma cells, loss of CD19^+^ cycling precursors such as plasmablasts and ABCs, and the intrinsically short lifespan of autoreactive plasma cells in *Tlr7*^GoF^ mice. In contrast to CD20-targeting antibody depletion therapy (Khodadadi et al. 2015), B cells were completely absent from the spleen post-anti-CD19 CAR-T cell administration. The effectiveness of CAR-T therapy in eliminating tissue-resident B cells points to its superior tissue penetration.

Our findings have established that the persistence of the autoantibody-secreting cell cohort is driven by continuous replenishment from proliferative precursors rather than by long-lived plasma cells. These precursors likely include both plasmablasts and less differentiated B cell populations such as ABCs. While we show that plasmablasts and their progeny sustain autoantibody production over extended periods, the more rapid decline of antibodies derived from cells labelled by *Jchain^ERT2cre^* compared to *Cd79a^ERT2cre^* suggests that *Jchain*-negative B cells — i.e., plasmablast precursors — also contribute to maintaining the long-lived autoantibody response. This interpretation is further supported by the rapid recurrence of autoantibodies following αBCMA CAR-T cells therapy, which included autoantibodies produced by cells labelled prior to treatment. Together, these results indicate that BCMA-negative precursors sustain the autoantibody response over time and underlie disease relapse.

Our single-cell analysis revealed a subset of proliferating ABCs that are transcriptomically related to plasmablasts, although their close proximity in the UMAP may partly reflect shared expression of proliferation-associated genes. These cycling ABCs — and any other self-renewing precursors — may continuously seed the plasma cell compartment. Indeed, we identified clonal relationships between ABCs and plasma cells in both the spleen and bone marrow, adding to the evidence that ABCs can serve as precursors of autoreactive plasma cells (Nickerson et al. 2023).

Exposure to nucleic acid-associated antigens may provide continuous stimulation that maintains the cycling of autoreactive B cells. Lupus-prone mice exhibit abundant extramedullary haematopoiesis in the spleen, creating an environment enriched in dying cells, extruded erythroblast nuclei, and mitochondria which are all potential TLR7 ligands (Caielli, Wan, and Pascual 2023).

In our working model, chronic TLR7 signalling sustains a proliferative pool of ABCs and potentially other precursors populations that continuously replenish the plasma cell compartment, independently of the generation of new autoreactive B cells. We were nevertheless surprised to find that plasma cells were uniformly short-lived across all tissue sites, as supported by the rapid loss of *Prdm1^ERT2cre^* tag-switched antibodies and the near-complete depletion of serum autoantibodies within seven days of CD19-CAR-T cell therapy. This contrasts with previous mouse studies suggesting that autoreactive plasma cells are often long-lived. The discrepancy may reflect limitations of BrdU labelling, where BrdU-low plasma cells arising from slowly dividing precursors can be misinterpreted as true long-lived, non-proliferating plasma cells. While the half-life of IgG in mice is approximately 21 days, the rapid depletion of serum autoantibodies within seven days of CAR-T therapy, and the persistence of immune complexes in the spleen 35 days post-treatment, suggests that FcRn saturation by these complexes may accelerate IgG degradation (Pyzik et al. 2023).

The short lifespan of plasma cells in *Tlr7*^GoF^ mice is unlikely to result from their preferential accumulation in the spleen. Although the bone marrow is typically a better survival niche and bone marrow displacement causes rapid plasma cell death (Nutt et al. 2015), the spleen can support long-lived plasma cells under inflammatory conditions (Khodadadi et al. 2019). Yet, unlike WT plasmablasts, which frequently colocalise with CD11c^hi^ dendritic cells (García De Vinuesa et al. 1999) producing APRIL and BAFF — factors that are critical for plasma cell longevity (Schuh, Mielenz, and Jäck 2020) —*Tlr7*^GoF^ plasma cells were typically found in areas lacking CD11c^hi^ dendritic cells. Additional extrinsic and intrinsic factors, including chronic TLR7 signalling, BCR crosslinking, cytotoxic T cell-mediated killing, and FcγIIB crosslinking by immune complexes, may also accelerate plasma cell death.

Unexpectedly, we found that the spleen harboured the highest proportion of self-reactive antibody-secreting cells relative to cellularity compared to the bone marrow and other inflamed tissues. In activated B cells, increased TLR7 signalling promotes the expression of the chemokine receptor CXCR3 (Brown et al. 2022), which likely contributes to the retention of ABCs within the inflamed spleen. Consistent with this, we observed a reduced frequency of IgG2c^+^ plasma cells in the bone marrow compared to the spleen. Both class switching to IgG2c and upregulation of CXCR3 are hallmark features of ABCs and are regulated by the transcription factor T-bet (Johnson et al. 2020).

Analysis of spleens from patients with SLE and associated conditions also revealed abundant plasma cells in this organ. Surprisingly, plasma cells were found to concentrate around splenic arterioles and other smaller vessels, suggesting the presence of plasma cell-attracting chemokines and/or growth factors at these sites. The close proximity of plasma cells with vascular endothelium suggests subendothelial deposition of autoantibodies, which may contribute to endothelial damage, release of DAMPs including nucleic acids, and further type I-IFN production.

The finding that αBCMA CAR-T therapy insufficiently depleted autoantibody-secreting cell precursors supports the conclusion that therapies targeting all CD19^+^ cells may be more effective than those solely focused on plasma cells. Anti-BCMA therapies are now being used to treat relapses post-CD19 CAR-T cell therapy (Müller et al. 2025). While we do not exclude the possibility of long-lived plasma cells in some lupus patients and in patients with other systemic autoimmune diseases, particularly when germinal centres give rise to pathogenic autoreactive plasma cells, our findings suggest that pan-B cell-targeting strategies may offer more sustained remission.

Finally, our finding that newly produced self-reactive B cells are continuously recruited into the autoantibody response, and that these cells rapidly drive relapse in *Tlr7*^GoF^ mice treated after αCD19 CAR-T therapy, indicate that increased TLR7 signalling profoundly impairs negative selection of self-reactive B cells. Consequently, patients with strong genetic predisposition are unlikely to maintain a durable B cell reset following CAR-T cell therapy and will be prone to relapse. Therapies that limit maturation and/or terminal differentiation of self-reactive B cells into ABCs and autoantibody-secreting cells — e.g., blocking B-cell activating factor (BAFF) or TLR7 signals — may help sustain the CAR-T–induced reset if given prior to B cell re-emergence.

## Limitations of the study

Limitations of this study include species-specific differences in TLR7 and TLR8 signalling and tissue microenvironments that may not fully model all manifestations of human lupus. For example, plasma cell lifespan may be unusually short in the presence of strong TLR7 signalling, potentially affecting outcomes observed with CAR-T cell depletion. Additionally, *Tlr7*^GoF^ (Y264H) mice used here do not develop severe nephritis, which in humans could represent another tissue niche for plasma cell accumulation. Lastly, the CAR-T construct used in this study was designed for mice, and CAR-T constructs used in human settings may differ in their depletion efficiency. This study focuses on monogenic lupus which is a small fraction of human lupus cases that typically presents in childhood. Adult-onset disease is generally associated with milder genetic predisposition and stronger environmental contributions, and thus relapses after CAR-T cell therapy are likely to be rarer and occur later after waning of CAR-T cell activity.

## Methods

### Mouse models

*Tlr7*^GoF^ (Y264H) and *Dnase1L3^-/-^* mice were generated at the Australian National University (ANU), Canberra, Australia. *Tlr7*^GoF^ mice were created as described (Brown et al. 2022) in a C57BL/6NCrL background and were backcrossed at the Francis Crick Institute to C57BL/6J. *Dnase1l3^-/-^* mice were created at the ANU on a C57BL/6NCrl background by targeting Exon5 by direct pronuclear injection of CRISPR/Cas9 complex with crRNA guides. A 74bp deletion was induced at exon5 generating a full knockout of Dnase1L3. They were rederived onto a C57BL/6J background at the Francis Crick Institute. *Trex1^-/-^* mice were from LRI CRUK (Morita et al. 2004).

*Igk^Tag^* mice were a kind gift from from G. Victora (Schiepers et al. 2023). *Cd79a^ERT2cre^* (Mb1-ERT2cre) (Hobeika et al. 2015) and *ROSA26-mTmG* mice (Muzumdar et al. 2007) were obtained from The Jackson Laboratory (Stock #033026 and 007676). *Prdm1^ERT2cre^* mice were a kind gift from Elizabeth Robertson (Elias et al. 2017). *Jchain^ERT2cre^* mice were gifted by Dinis Calado (Xu, Barbosa, and Calado 2020).

All mice were held at the Biological Research Facility at the Francis Crick Institute under specific-pathogen-free conditions. All mouse procedures were approved by the Francis Crick Institute’s strategic oversight committee (Animal Welfare and Ethical Review) and by the Home Office, UK. All animal care and procedures followed guidelines of the UK Home Office according to the Animals (Scientific Procedures) Act 1986. The age of the mice ranged from 4 – 35 weeks as specified in figure legends.

### ELISA

96-well High binding ELISA Microplates (Greiner) were coated with poly-L-lysine (Sigma-Aldrich) diluted 1:50 in dH2O and incubated for 4+ hours at room temperature. To determine titres of both Flag- and Strep-tag carrying serum antibodies, antigen coating was performed by coating 10 columns (column 1-5 and 7-12) with 50µg/mL DNA (Sigma-Aldrich, D8661), 1µg/mL Ro60 (pro-329-b, PROSEC), 1µg/mL sm/RNP (AROTEC, ATR01), or 2µg/mL Goat Anti-Mouse Kappa-UNLB (SouthernBiotech), and coating the two middle columns with 10µg/mL CGG (Antibodies Online) in Carbonate/Bicarbonate buffer (pH 9.6) at 4°C overnight. Plates were washed 4 times with 0.05% Tween-PBS on a BioTek ELx405 Select CW plate washer and blocked with 1% BSA in PBS for 2 hours at room temperature. Mouse serum was diluted 1:100 (anti-DNA and anti-IgK detection) or 1:50 (anti-Ro60 and anti-sm/RNP detection) in blocking buffer and applied in duplicates on opposite sides of the plate. Monoclonal mouse IgG CGG-specific antibodies carrying C_κ_ chains with a Flag- or Strep-tag identical to those produced by *Igk^Tag^* mice (Schiepers et al. 2023) were used as standard controls to verify similar detection of Flag- and Strep-tag antibodies. Flag- and Strep-tag monoclonal antibodies were serially diluted (starting concentration 0.20µg/mL) at a 1:3 ratio and added to the two middle columns of the plate. Plates were incubated with serum and monoclonal antibody standards at 4°C overnight. After washing with 0.05% Tween-PBS, the 6 left columns were incubated with anti-Flag-tag (FG4R, Invitrogen) and the 6 right columns with anti-Strep-tag (5A9F9, Genscript) HRP-conjugated detection antibodies for 1 hour at room temperature. Detection antibody was diluted in blocking buffer and dilutions were calibrated so that the standard curves generated by Flag- and Strep-tag monoclonal antibodies were equal. Plates were washed with Tween-PBS and incubated with 3,3′,5,5′-tetramethylbenzidine (TMB) substrate (slow kinetic form, Sigma-Aldrich) and the reaction was stopped after 25min with 1N HCl. Optical density (OD) absorbance was measured at 450 nm and normalized to background absorbance at 570 nm using a Safire^2^ or an Infinite M1000 Tecan plate reader. The average of duplicate values in Flag- and Strep-tag fractions were calculated.

### Generation of recombinant antibodies

CGG-specific Flag- and Strep-tag monoclonal antibodies were produced by transfecting HEK293T cells with heavy-chain plasmid together with either Flag- or Strep-tag light-chain plasmids. Plasmids were a kind gift from G. Victora (Schiepers et al. 2023). Plasmids were amplified and isolated from culture using a Midi kit for endotoxin-free plasmid DNA (NucleoBond Xtra Midi Plus EF, Macherey-Nagel) following manufacturer’s instructions. HEK293T cells were transfected in serum-free media using lipofectamine 2000 Transfection reagent (Thermo Fisher) following manufacturer’s instruction. Antibodies were purified from the supernatant after 2- and 7-days incubation using protein-G affinity chromatography. The concentration of aliquoted monoclonal antibody was determined by nanodrop before use in each ELISA assay.

### Tamoxifen induction and serum collection

In ERT2cre-expressing mouse lines the antibody immune response was time-stamped by administering 250 µL tamoxifen (APExBIO) dissolved in corn oil by oral gavage. For all experiments monitoring the autoantibody immune response in untreated mice, tamoxifen was given once at 60mg/mL or twice at 30mg/mL on day -2 and repeated on day 0. For CAR T cell depletion experiments, tamoxifen was given 6 to 9 days before CAR T cell administration and repeated two days after. To monitor autoantibody secretion from Flag^+^ and Strep^+^ cells, serum was collected on the day of the first tamoxifen administration, one week after tamoxifen, and again at 2–3-week intervals after that as indicated in the figures. All mice were bled from the saphenous vein into heparin-coated blood-collection tubes. Serum was separated by centrifugation at 1000 x g for 10minutes at 4°C, aliquoted, and stored at -20°C.

### ELISPOT

96 well MSIP (multiscreen, Merck) plates were pre-wet with 50μL 35% ethanol per well and incubated for 1 minute before washing four times with distilled water. 15μg/mL Goat Anti-Mouse Kappa-UNLB antibody (SouthernBiotech) or 100 μg/mL DNA (Calf thymus, D8661 Sigma-Aldrich) in PBS was added to the plate and incubated overnight at 4°C. Salivary/submandibular lymph nodes, Peyer’s patches, and spleen were isolated from the mice and single cell suspensions created by passing the organs through a 70μm filter. Bone marrow cells were isolated by removing one femur and tibia bone per mouse and flushing with PBS. Collected bone marrow suspension was filtered through a 40μm filter. All cell suspensions were washed in RPMI, resuspended in R10 medium and added to the ELISPOT plate at 3000 and 6000 (anti-mouse Kappa coated) or 200,000 and 400,000 (DNA coated) cells per well. Plates were covered with tinfoil to minimise evaporation and incubated overnight (16-18 hours) in a 37°C 5% CO_2_ incubator. For detection of antibody-secreting cells (ASCs) cells were first removed by washing thoroughly four times with PBS. HRP-conjugated detection antibodies specific to mouse-IgG (abcam), mouse-IgA (abcam), Flag-tag (L5, Biolegend), or Strep-tag (5A9F9, Genscript) were added at 0.5-1μg/mL and plates were incubated for 2 hours at room temperature. After washing with PBS, 50μL of TMB ELISpot substrate (Mabtech) was added per well. Colour development was monitored and the reaction stopped after approximately ten minutes by washing extensively with deionized water. Plates were allowed to dry overnight at 4°C and ASCs were quantified using a CTL ImmunoSpot S5 spot analyser. The average number of spots of two replicates was calculated.

### Immunofluorescence

Spleens were isolated from mice and directly fixed for 24hrs in 10%NBF before processing to wax using a Tissue-Tek VIP® 6 AI processor. 3 µm FFPE sections were cut and baked for 1hr at 60°C before staining was performed on the Leica Bond Rx platform. Hydrogen peroxide (3%) was used to block the endogenous peroxidase and 0.1% BSA solution was used for protein blocking. Antigen retrieval stripping steps between each antibody were performed with Epitope Retrieval Solution 2 for 20 minutes. Triple immunofluorescence staining was performed and antibodies were applied with OpalÔ pairings in the following order: Figure 5 and Figure 7F: anti-Ig (ThermoFisher, A-10666) at 1:500 with Opal 570 at 1:500, anti-CD138 (R&D Biotechne, AF3190) at 1:400 with Opal 520 at 1:500 and anti-CD19 (abcam, ab245235) at 1:200 with Opal 690 at 1:200. Consecutive slides were stained with anti-CD11c (CST, 97585) at 1:200 with Opal 520 at 1:500, anti-CD138 (R&D Biotechne, AF3190) at 1:400 with Opal 570 at 1:500 and anti-CD19 (abcam, ab245235) at 1:200 with Opal 690 at 1:200. Figure S7: anti-CD11c (CST, 97585) at 1:200 with Opal 520 at 1:500, anti-CD3 (ab134096) at 1:500 with Opal 570 at 1:500, anti-CD19 (abcam, ab245235) at 1:200 with Opal 690 at 1:200.

Bond anti-rabbit Polymer (Leica, RE7260-CE) was used as secondary detection for antibodies raised in rabbit. ImmPRESS® HRP Horse Anti-Goat IgG Polymer Detection Kit (Vector Laboratories, MP-7405) was used as secondary for antibodies raised in goat. Slides were counterstained with DAPI (Thermo Scientific, 62248) 1:2500. Slides were mounted with Prolong Gold Antifade reagent (Invitrogen, P36934). Images were acquired using an Evident/Olympus VS200 slide scanner with a UPlanXApo 20x/0.8 objective.

### Flow Cytometry

Spleens were isolated, weighed, and passed through a 70µm filter to create single cell suspensions. Bone marrow cells were isolated from one femur and one tibia bone per mouse by flushing with PBS. Cell suspensions were treated with 0.5 mL Red-Blood-Cell Lysis buffer for five (bone marrow) or ten (spleen) minutes at room temperature before washing with 10mL FACS buffer (1% FCS, 50mM EDTA in PBS). 10 × 10^6^ cells were blocked with 1µL Fc-blocking antibody (TruStain FcX, Biolegend) for 10min on ice, and subsequently stained with antibody cocktails in 100 µL FACS buffer. For a list of antibodies used see Extended Materials. Samples were stained for 30 min on ice in dark conditions and washed with 2mL FACS buffer before flow cytometry acquisition on a Fortessa X-20 instrument. Flow cytometry data were analysed using Flowjo (v.10.6.2).

For sorting of plasma cells and ABCs for single cell RNA sequencing, spleen and bone marrow cell suspensions from three mice total were prepared similarly to described above. Single cell sorting into 384 well plates prepared with lysis buffer (see FLASH-seq description) was calibrated on a BD FACSAria III Cell Sorter. Single cells were sorted into each well with the exception of two wells kept for negative controls. A 100μm nozzle was used to minimise shear stress. After ended sort, plates were covered with aluminium foil seals, briefly centrifuged and then placed on dry ice before being stored at -80°C until library preparation.

### Retroviral vector design and production

Retroviral transfer plasmids were constructed on a SFG backbone using standard molecular biology techniques. The transfer genome for each retroviral vector contained a CAR consisting of scFv fragments derived from anti-mouse CD19 1D3, anti-mouse BCMA 30H1C6, or anti-human IL13RA-2 06B5, linked in frame to the same CD8STK-CD8TMD-41bbz signalling domains.

For retroviral vector production, Phoenix-Eco packaging cells (ATCC CRL-3214) were cultured in IMDM (Gibco) supplemented with 10% heat-inactivated FCS (Sigma) and 1% penicillin-streptomycin (Gibco). 2 × 10⁶ cells were plated in 100 mm tissue culture dishes 24 hours prior to transfection. At 60–70% confluency, transfection was performed using GeneJuice (Merck Millipore) according to manufacturer guidelines: Briefly, 30 µL GeneJuice was incubated with 470 µL serum-free IMDM for 5 minutes, then combined with 2.6 µg pCL-Eco packaging plasmid (Addgene #12371) and 4.7 µg of the transgene vector. The DNA-GeneJuice complex was incubated for 20 minutes at room temperature before dropwise addition to cells. After 24 hours, medium was replaced with 5 mL fresh IMDM (Gibco) containing 10% FCS. Viral supernatants were harvested at 48-and 72-hours post-transfection, pooled, and centrifuged (400 xg, 5 minutes) to remove cellular debris. Supernatants were filtered through 0.45 µm PES syringe filters (Millipore) and snap-frozen in an ethanol/dry ice bath before storage at −80°C.

### Primary Mouse T Cell Isolation

Spleens from C57BL/6J mice (6–8 weeks) were aseptically dissociated through 70 µm cell strainers (Corning) into PBS. Red blood cells were lysed using 2 mL ACK lysis buffer (Gibco) for 5 minutes at room temperature, neutralized with 10 volumes PBS, and filtered through 40 µm strainers. Cells were washed twice in PBS (400 xg, 5 minutes) and resuspended in complete mouse T cell medium: RPMI-1640 supplemented with 10% FBS, 1xglutaMAX (Gibco), 10 mM HEPES (Sigma), 1 mM sodium pyruvate (Gibco), 1× non-essential amino acids (Gibco), 50 µM 2-mercaptoethanol (Gibco), and 1% penicillin-streptomycin (Gibco). For downstream transduction, isolated T cells were activated by culturing with 5 µg/mL concanavalin A (ConA; Sigma) for 24 hours prior to retroviral transduction.

### Retroviral transduction of primary mouse T cells

Non-tissue culture 6-well plates (Greiner Bio-One) were coated with 2 mL RetroNectin (Takara Bio, 10 µg/mL in PBS) at 4°C for ≥24 hours. Coated wells were blocked with PBS containing 2% BSA (Sigma) for 30 minutes at room temperature, followed by two PBS washes. Activated splenocytes were harvested and resuspended at 1 × 10⁶ cells/mL in complete mouse T cell medium. 1 mL of T cells were combined with 3 mL of viral supernatant, and added to RetroNectin-coated wells. Plates were spinoculated (1000 xg, 32°C, 90 minutes) then incubated overnight at 37°C/5% CO₂. Transduced cells were washed twice with PBS, resuspended in complete medium containing 50 IU/mL recombinant IL-2 (Merck Millipore). Two days later, transduction efficiency was assessed by flow cytometry by staining cells for Thy1.1 which is expressed on the CAR T construct (OX-7, BioLegend). Transduction efficiencies ranged from 57 to 95%. T cells were frozen and stored at -80°C or in liquid nitrogen prior to downstream application.

### CAR T cell depletion

To fate-map the B cell immune response Igĸ-tag switching was induced in *Tlr7*^GoF^ *Cd79a^ERT2cre^ Igk^Tag/Tag^* mice by tamoxifen administration (as described above). 3-6 days after the last tamoxifen treatment, mice received a single dose of 0.2mg/g cyclophosphamide (ie. 5mg per 25g mouse weight) intraperitoneally. One day after (day 0), mice received 1.6×10^6^ CAR T cells I.V.

B cell depletion and CAR T cell persistence was monitored up to 140 days after CAR T cell treatment by flow cytometry: 30µL blood was stained and prepared using eBioscience 1-step Fix/Lyse Solution according the manufacturer’s protocol. Samples were stained in 50µL volume using antibodies: anti-B220-AlexaFluor488 (RA3-6B2, Biolegend), anti-CD19-APC (6D5, Biolegend), anti-CD3-AlexaFluor700 (17A2, Biolegend), anti-Thy1.1-PE (OX-7, Biolegend) at 1:200 dilution, and Live/Dead-Fixable Near IR (Invitrogen) at 1:1000 dilution. For enumeration of total B and T cells, 10µL Precision Count Beads (Biolegend) were vortexed and added to each sample before flow cytometry acquisition. Serum was isolated from leftover blood by centrifugation at 1000xg for 10minutes at 4°C, aliquoted, and stored at -20°C.

### Single cell FLASH-sequencing

Cells were sorted into 384-well plates prepared with 1µl FLASH-seq lysis buffer, and processed as per the published low amplification protocol (Hahaut et al. 2022) with minor adjustments. Briefly, reverse transcription was performed in a 5µl reaction volume with 15 cycles of pre-amplification. cDNA was then diluted 1:7 with nuclease-free water, and 0.8µl was taken forward into the tagmentation reaction, with 1µl 2X TD and 0.2µl ATM (FC-131-1096, Illumina). Low-volume pipetting was carried out using the SPT Labtech Mosquito liquid handling platform. Libraries were amplified with IDT Nextera UDI oligonucleotides using 14 cycles of amplification; equi-volume pooled, and cleaned up using a double-sided size selection with SPRI Select. The pool was quantified by Qubit High Sensitivity dsDNA assay, sized using Agilent TapeStation 4200 D1000 assay, and diluted to 4nM for sequencing on Illumina NovaSeq 6000.

### Data analysis

Raw demultiplexed fastq files were processed using the nf-core rnaseq pipeline (v3.17.0) (Patel et al. 2025). Star-rsem was used to align reads to the GRCm38 (mm10) genome and create count tables.

### Single cell analysis

All bioinformatics analyses were performed in the R studio software (version 4.4.3) and graphs were made using ggplot2. To create count tables of gene expression for each plate, the count matrices were converted to Seurat objects using R package Seurat (v. 5.3.0). Cells with less than 200 genes and features present in less than 3 cells were excluded. Mitochondrial content was negligible across each object. Metadata information about cell type, fluorescent marker and mouse of origin was added before the Seurat objects from all five plates were merged. The merged dataset was normalised using the SCTransform function and Uniform Manifold Approximation and Projection (UMAP) was used to reduce dimensions

### BCR alignment and clonal analysis

VDJ alignment for each sample was performed using BraCeR (Lindeman et al. 2018). Clonality plots were created using BraCeR, which groups clonally related cells into clonotypes based on the V-and J-gene assignments of reconstructed chains in each cell. Clonal trees were generated using R packages dowser (Hoehn, Pybus, and Kleinstein 2022) and ggtree (Yu 2022) as part of the Immcantation tool suite.

### Human samples and ethics

Human spleen samples were obtained post-splenectomy for clinical reasons indicated in Supplemental Table 1. Samples were formalin-fixed and paraffin-embedded prior to sectioning and immunofluorescence staining. The work was approved by Istanbul Faculty of Medicine Clinical Research Ethics Committee and by the Francis Crick Institute Human Biology Facility. These patients were managed before biological therapies were routinely available.

### Statistics

Statistical analysis was carried out using GraphPad Prism v.10.0. ELISPOT quantification data was analysed using one-way ANOVA, followed by Tukey’s multiple comparison testing. Where two genotypes were compared side-by-side two-way ANOVA followed by Tukey’s multiple comparisons testing was used. For survival curves a simple survival analysis (Kaplan-Meyer) was used and the Logrank (Mantel-Cox) test p value summary reported. Differences in the frequency of Flag^+^ expression among different cell types (Supplemental Figure 1) was tested by Friedman tests with Dunn’s multiple comparisons correction. FACS data, spleen mass and autoantibody titres among four genotypes (Supplemental Figure 2) were analysed by one-way ANOVA followed by Tukey’s multiple comparison testing. Data showing the frequency of GFP^+^ cells among two genotypes (Supplemental Figure 3) was analysed using two-way ANOVA followed by Sidak’s multiple comparisons test. Mouse spleen mass data (Figure 7), FACS data on plasma cell populations (Supplemental Figure 2), and data on recruitment indices (Supplemental Figure 4) between two genotypes were analysed using unpaired t-test. ELISPOT comparison between spleen and bone marrow cells was analysed using paired t-test (Figure 4). Data were filed using Microsoft Excel 2024 and graphed using GraphPad Prism or using R studio software (v. 4.4.3) and graphed using ggplot2.

## Acknowledgements

We thank The Francis Crick Institute: Experimental Histopathology, E. Nye, A. Mikolajczak and team for their excellent work on multiplex immunofluorescent staining of spleen sections; Genomics, R. Goldstone, D. Snell, M. Rodriguez and team for their invaluable support in the development and optimisation of the FLASH-sequencing protocol; Bioinformatics & Biostatistics, P. East, M. Llorian Sopena, J. Campbell for vital help with bioinformatics pipelines; Biological Research Facility, N. Chisholm, J. Murphy, S. Kuncova, A. Townsend, L. DeRosa, for their excellent care-taking of animal models, and D. Gibbins for her exceptional help in colony management; Flow Cytometry, A. Riddell, A. Agua-Doce, and team for support with flow cytometry and sorting experiments; Cell Science and Structural Biology, T. Cooper, and A. Borg for help with the production of monoclonal antibodies.

This work was enabled by support from a Wellcome Trust Discovery Award (302393/Z/23/Z) and from the Francis Crick Institute (CC2228), which receives its core funding from Cancer Research UK, the UK Medical Research Council and the Wellcome Trust. C.G.V. has also received funding from a Royal Society Wolfson Fellowship. A.G. was supported by a MSCA/ UKRI Post-doctoral Fellowship (EP/Y031091/1) and an EMBO post-doctoral Fellowship (EMBO ALTF 654-2022). A. R-R. was funded by Ministerio de Ciencia e Innovación (PRE2020-091873).

## Author contributions

A.G. and C.G.V. conceptualised the study and wrote the manuscript with contribution from all authors. *In vivo* experimental work was led by A.G. with assistance from H.W. J.W. transduced CAR-T cells for use in *in vivo* work under supervision by P.M. and L.L. who also contributed conceptually. FLASH-sequencing was performed by D.M.S. Bioinformatics analysis was performed by A.G. with assistance from A.R-R. Human spleen samples and associated information were provided by E.G., G.Y., led by B.A.E. Multiplex immunofluorescence was performed by A.M. A.R. contributed important conceptual input and discussions.

