## Supplemental Figures and legends for "Autoantibody origins in lupus and in relapse post CAR-T therapy"

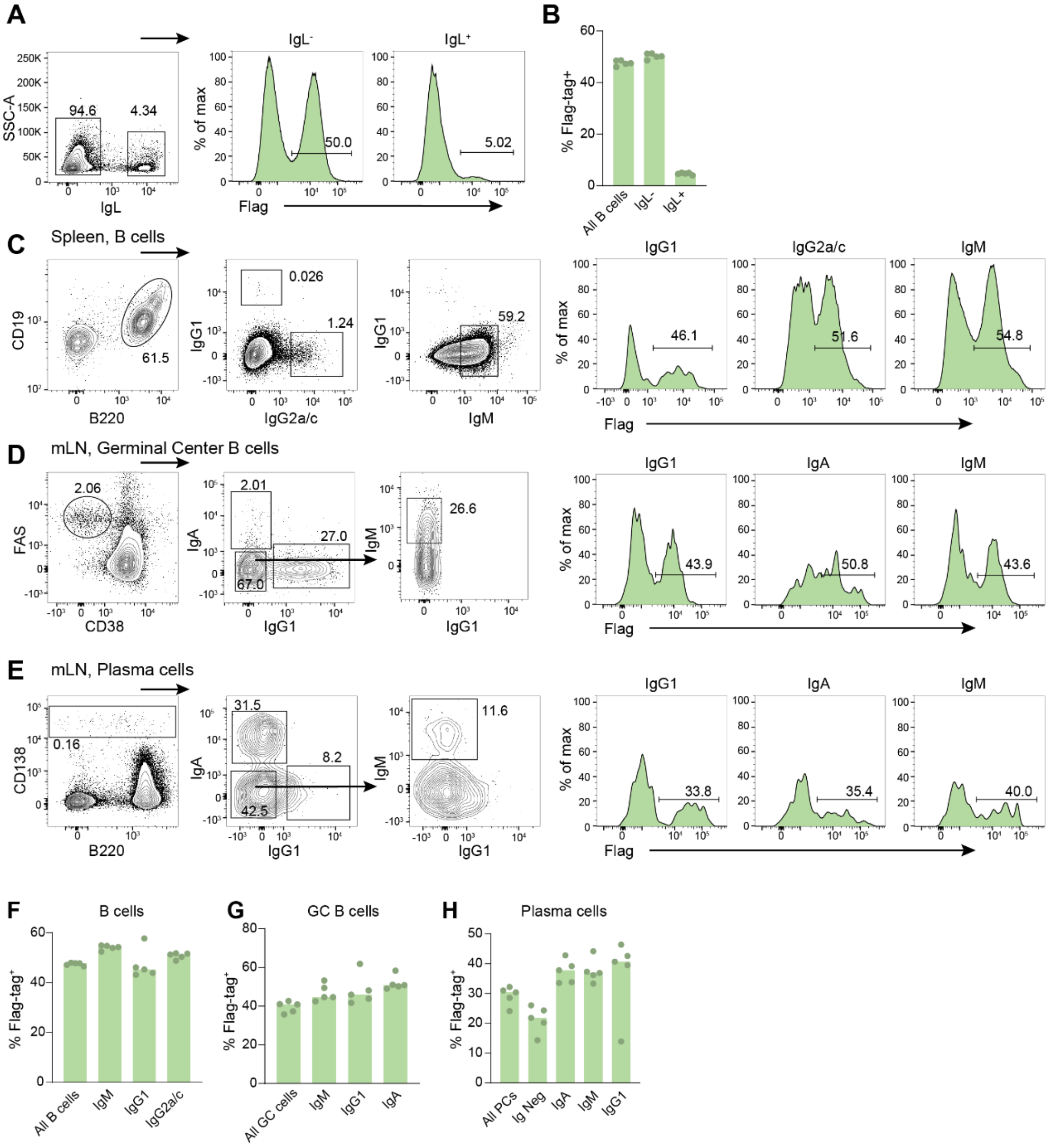


**Supplemental Figure S1: Igκ-tag detection in isotype switched cells. (A)** Flow cytometry plot showing Igl detection in splenic CD19+ B220+ B cells (left) and Igκ-tag detection in Igl- and Igl+ B cells (right) from a *Igk*$^{WT/Tag}$ mouse. **(B)** Quantification of Igκ-tag detection from A. **(C-E)** Flow cytometry gating strategy of isotype-switched cells and Igκ-tag detection in each population in total splenic B cells (C), mesenteric lymph node germinal centre B cells (D), and mesenteric lymph node plasma cells (E). **(F-H)** Quantification of Igκ-tag detection of data in C-E. Dots represent individual values, column height represents median value.

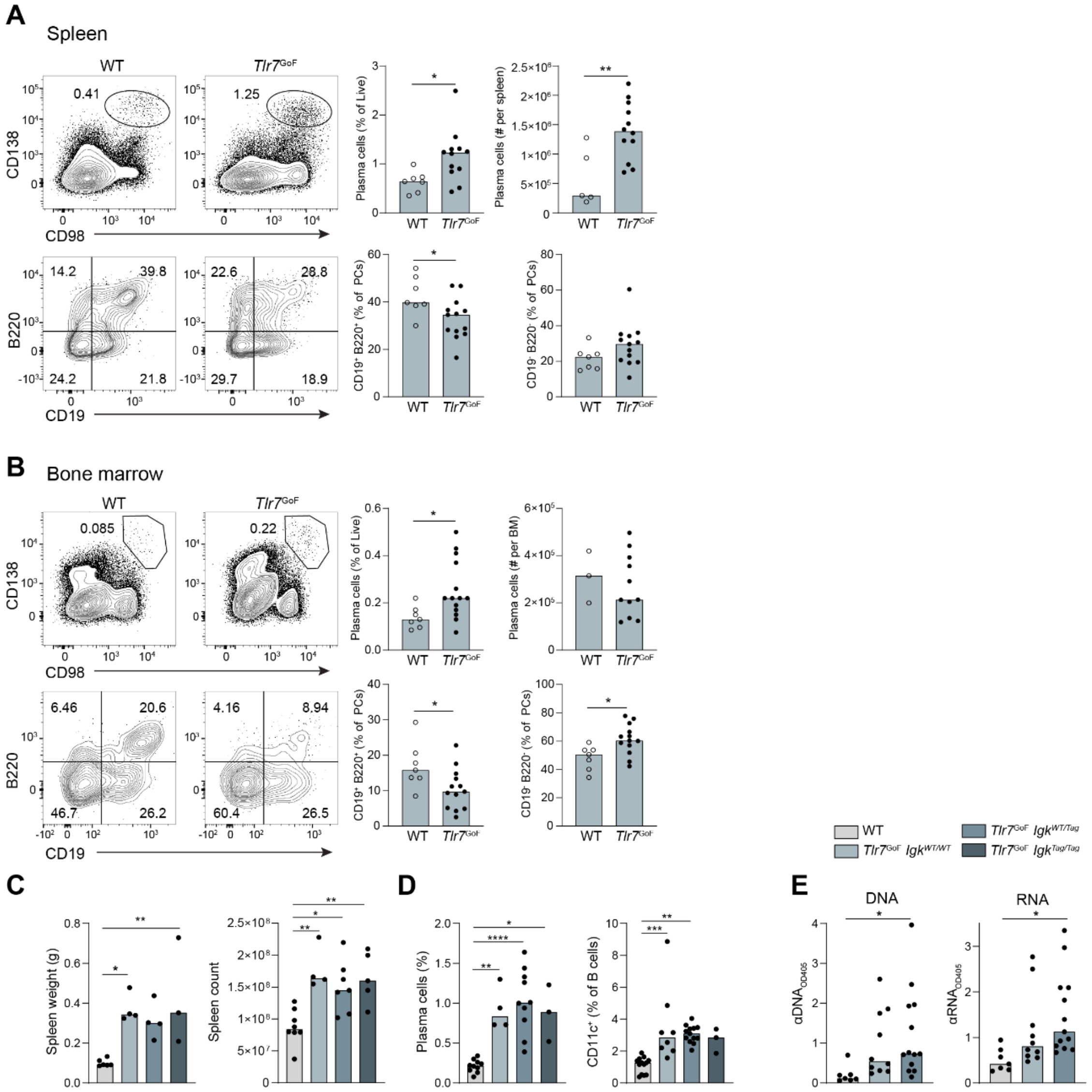


**Supplemental Figure S2: Plasma cell populations and lupus phenotyping in *Tlr7*$^{GoF}$ *Igk*$^{Tag}$ mice.** **(A-B)** Flow cytometry plots (left) and quantification (right) of plasma cells and expression of CD19 and B220 in spleen (A) and bone marrow (BM) (B) of WT and *Tlr7*$^{GoF}$ mice. Pooled data from two experiments with 3-10 mice per group, bars show median values. **(C-E)** Analysis of spleen size and cellularity (C), frequency of plasma cell and CD11c$^{+}$ ABC populations (D) and anti-DNA and anti-RNA serum antibody titres (E) in WT and *Tlr7*$^{GoF}$ mice with 1-2 copies of the *Igk*$^{Tag}$ allele. Pooled data from 3 experiments. Dots show individual values, column height represents the median.

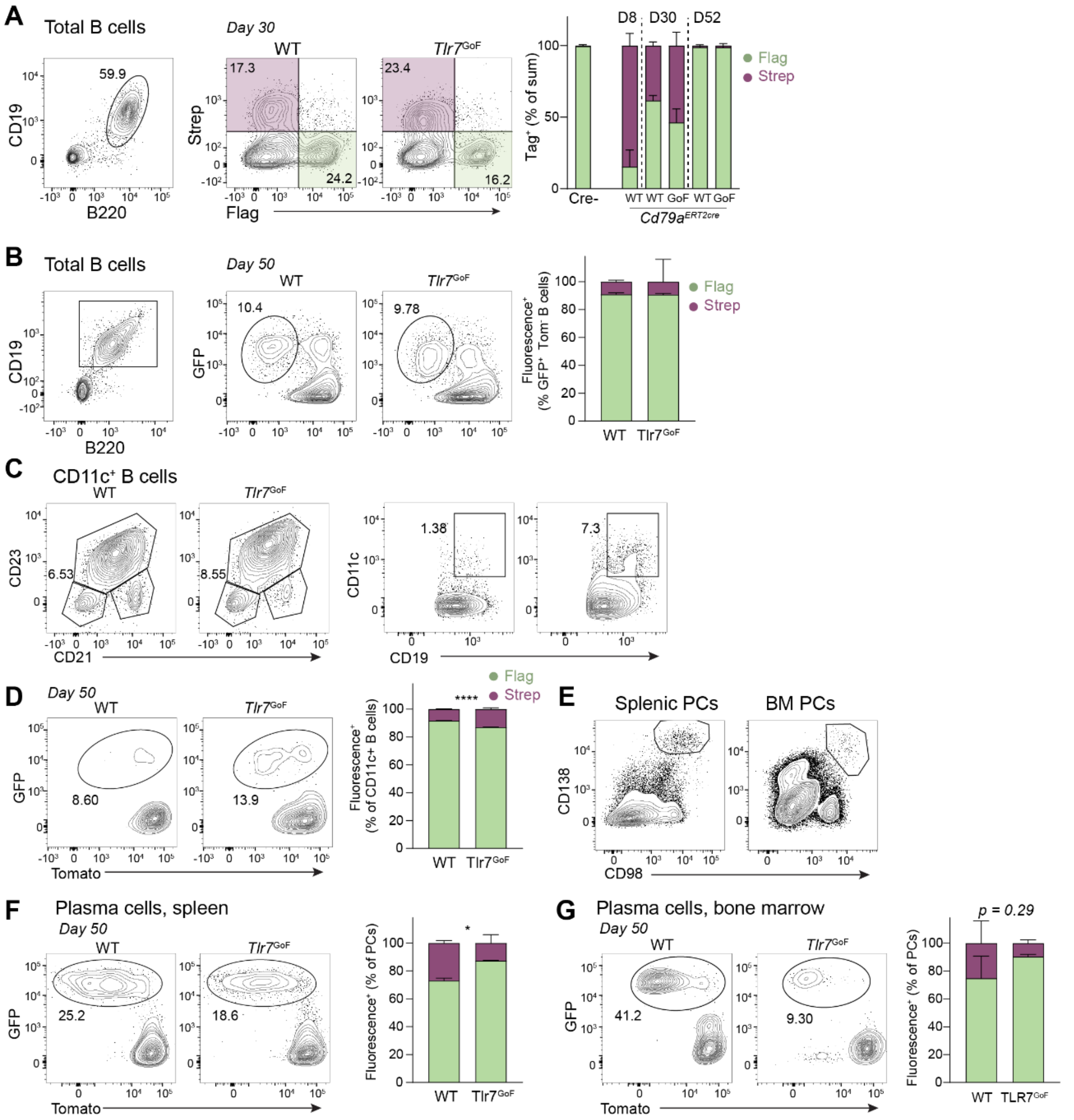


**Supplemental Figure S3: *Tlr7*$^{GoF}$ mice have slower turnover of CD11c$^+$ ABCs but faster turnover of plasma cells. (A)** Gating strategy of B cells (left) and surface expression of Flag- and Strep-tag (middle) in WT and *Tlr7*$^{GoF}$ *Cd79-Igk*$^{WT/Tag}$ mice 30 days after tamoxifen by flow cytometry. Relative contribution of Flag$^+$ and Strep$^+$ cells to total Igκ-tag$^+$ cells quantified over time (right). 2-5 mice per group, bars show median with 95% CI. **(B)** Flow cytometry gating of B cells and tdTomato/GFP expression in WT and *Tlr7*$^{GoF}$ *Cd79a-mTmG* mice 50 days after tamoxifen. **(C-D)** Gating of CD11c$^+$ ABCs (C) and tdTomato/GFP expression (D) in WT and *Tlr7*$^{GoF}$ *Cd79a-mTmG* mice. **(E-G)** Gating of plasma cells in spleen and bone marrow (BM) (E) and tdTomato/GFP expression in *Tlr7*$^{GoF}$ *Cd79-mTmG* mice (F-G). Relative fractions of tdTomato$^+$ and GFP$^+$ cells were quantified 50 days after tamoxifen. 2-4 mice per group; bars show median with 95% CI.

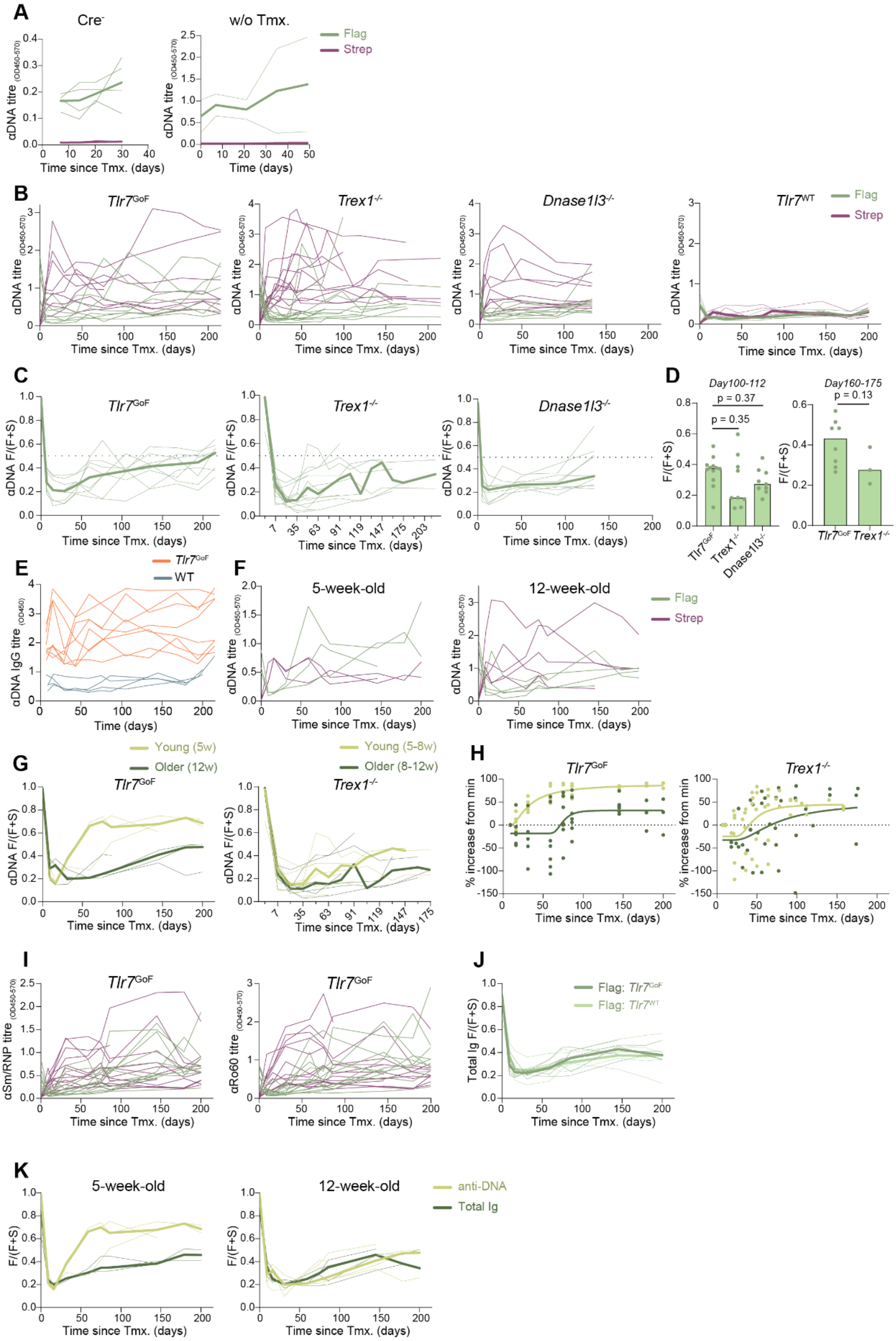


**Supplemental Figure S4: Autoantibody responses in *Tlr7*$^{GoF}$ and *Trex1*$^{-/-}$ mice. (A)** Anti-DNA antibody titres in tamoxifen treated *Igk*$^{Tag}$ *Cre*$^{-}$ mice (left) and *Tlr7*$^{GoF}$ *Cd79a-Igk*$^{Tag}$ mice without tamoxifen treatment (right) at consecutive timepoints. **(B)** Flag$^{+}$ and Strep$^{+}$ anti-DNA titres of individual mice corresponding to Figure 1G-H (left) and of *Tlr7*$^{WT}$ *CD79a-Igk*$^{Tag}$ mice (right). **(C)** Recruitment index of anti-DNA antibodies *(Flag / (Flag + Strep))* after tamoxifen treatment. *Tlr7*$^{GoF}$ *Cd79a-Igk*$^{Tag}$

mice (left), *Trex1*$^{-/-}$ *Cd79a-Igk*$^{Tag}$ mice (middle), and *Dnase1l3*$^{-/-}$ *Cd79a-Igk*$^{Tag}$ mice (right). **(D)** Recruitment index as in C at two time-intervals (100-112 days (left) and 160-175 days (right)) after tamoxifen. 3-12 mice per group analysed in two independent assays. Dots represent individual values; column height represents the median. **(E)** Anti-DNA antibody titres in WT and *Tlr7*$^{GoF}$ mice over time. **(F)** Flag$^{+}$ and Strep$^{+}$ anti-DNA antibody titres in 5- and 12-week-old *Tlr7*$^{GoF}$ *Cd79a-IgK*$^{Tag}$ mice following tamoxifen treatment, corresponding to Figure 1I. **(G-H)** Recruitment index of anti-DNA antibodies after tamoxifen (G), and the percentage increase in Flag$^{+}$ antibodies after day 14 post-tamoxifen (H), calculated at consecutive timepoints in young and older *Tlr7*$^{GoF}$ *Cd79a-Igk*$^{Tag}$mice (left) and *Trex1*$^{-/-}$ *Cd79a-Igk*$^{Ta}$ mice (right). *Tlr7*$^{GoF}$ mice 5 and 12 weeks old, and *Trex1*$^{-/-}$ mice were 5-8 and 8-12 weeks old at tamoxifen treatment. Plots show 3 *Tlr7*$^{GoF}$ mice per group shown (representative of 3 experiments) and 6 *Trex1*$^{-/-}$ mice per group pooled into 2-week bins. **(I)** Flag$^{+}$ and Strep$^{+}$ anti-sm/RNP and anti-Ro60 antibody titres in individual *Tlr7*$^{GoF}$ *Cd79a-Igk*$^{Tag}$ mice after tamoxifen treatment corresponding to Figure 1K. **(J)** Recruitment index of total (Igκ$^{+}$) antibodies in *Tlr7*$^{WT}$ *Cd79a-Igk*$^{Tag}$ (light green) and *Tlr7*$^{GoF}$ (dark green) mice after tamoxifen administration. **(K)** Comparison of the recruitment index between total (Igκ$^{+}$) immunoglobulin and anti-DNA antibody titres in young (5w at tamoxifen) and older (12w at tamoxifen) *Tlr7*$^{GoF}$ *Cd79a-Igk*$^{Tag}$ mice after tamoxifen treatment. 3-4 mice per group; representative of 3 experiments. In all antibody titre plots: thin lines represent individual values; thick lines represent the median.

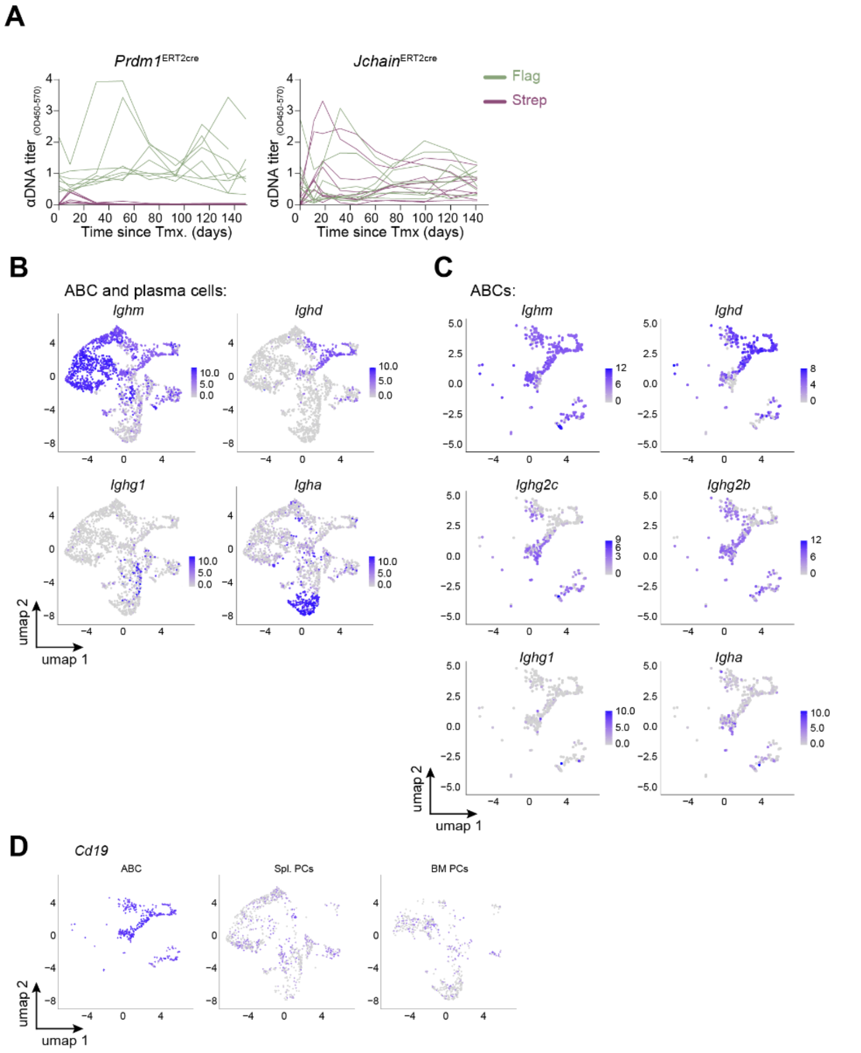


**Supplemental Figure S5: Isotype and Cd19 expression by single cell RNAseq. (A)** anti-DNA Flag$^+$ and Strep$^+$ antibody titres in individual *Tlr7*$^{GoF}$ *Prdm1*$^{ERT2cre}$ (left) and *Tlr7*$^{GoF}$ *Jchain*$^{ERT2cre}$ (right) mice after tamoxifen treatment as shown in Figure 2B and E. Thin lines represent individual mice, thick lines represent medians. **(B-C)** Expression of immunoglobulin isotype transcripts on umap from Figure 3B (B) and ABCs (C). **(D)** *Cd19* expression in ABCs and spleen- and bone marrow-resident plasma cells.

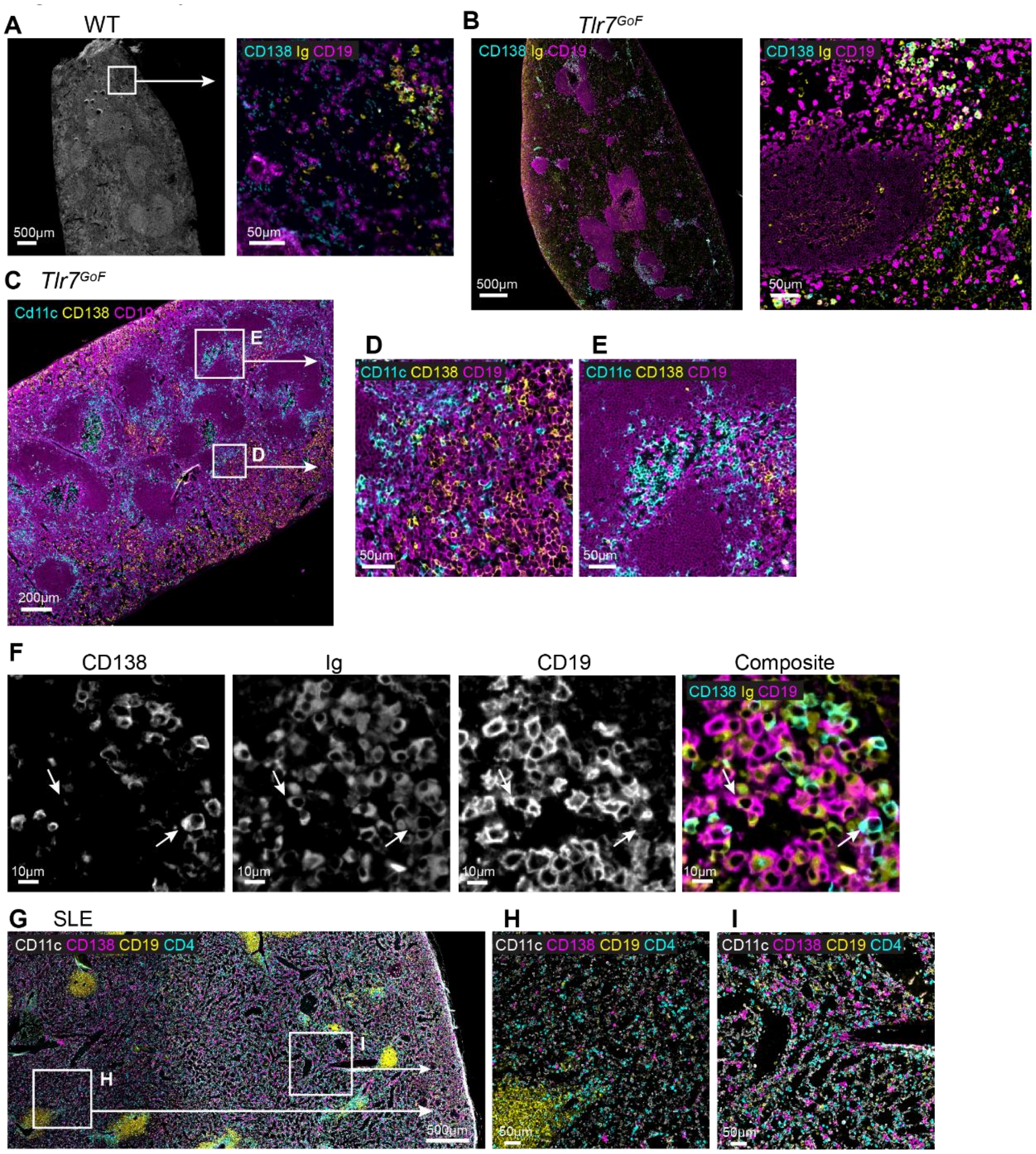


**Supplemental Figure S6: Plasma cell localization in *Tlr7*$^{\mathrm{GoF}}$ mouse and human spleen. (A)** Representative immunofluorescence images of WT mouse spleen. Left panel shows DAPI staining of a large spleen section; the white box indicates the red pulp area shown at higher magnification in the right panel. **(B)** Larger field of view of the spleen section shown in Figure 5B (left) and representative image showing plasma cells clustering near a B cell follicle (right). **(C-E)** Immunofluorescence images of a *Tlr7*$^{\mathrm{GoF}}$ mouse spleen stained for CD11c, CD138 and CD19. (C) Overview of a large spleen section; white boxes indicate the red pulp region shown in D and the B cell follicle region shown in E. **(F)** Consecutive immunofluorescence images showing single stains and a composite image of CD138, immunoglobulin (Ig), and CD19 staining in a plasma cell cluster from a representative *Tlr7*$^{\mathrm{GoF}}$ spleen. White arrows indicate a CD138$^-$ Ig$^+$ CD19$^+$ cell (left arrow) and a CD138$^+$ Ig$^+$ CD19$^-$ cell (right arrow). **(H-J)** Immunofluorescence images of a spleen section from a patient diagnosed with immune thrombocytopenic purpura (ITP), representative of findings observed in ITP, SLE, and control spleens. White boxes in H mark areas magnified in I and J.

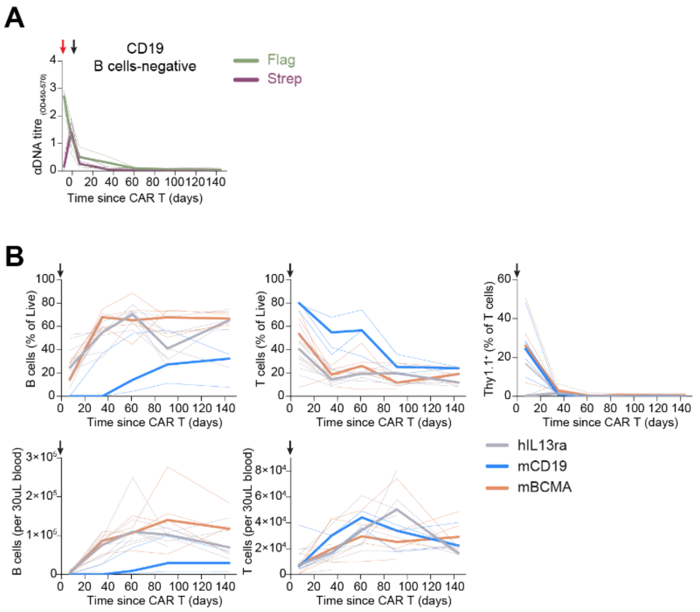


**Supplemental Figure S7: Monitoring after CAR-T cell therapy. (A)** Anti-DNA Flag$^{+}$ and Strep$^{+}$ antibody titres in four *Tlr7*$^{\mathrm{GoF}}$ *Cd79a-Igk*$^{Tag}$ mice aged 30 weeks at time of αCD19 CAR-T cell administration, in which B cells did not reconstitute. **(B)** Immunomonitoring of blood B cell, T cell and CAR$^{+}$ T cell (Thy1.1$^{+}$) populations after administration of αCD19, αBCMA, or αhIL13ra CAR-T cells. Thin lines represent individual mice, broad lines represent the median. Red arrow = Time of tamoxifen administration. Black arrows = time of CAR-T transfer.
